# Catch me if you can: Adaptation from standing genetic variation to a moving phenotypic optimum

**DOI:** 10.1101/015685

**Authors:** Sebastian Matuszewski, Joachim Hermisson, Michael Kopp

## Abstract

Adaptation lies at the heart of Darwinian evolution. Accordingly, numerous studies have tried to provide a formal framework for the description of the adaptive process. Out of these, two complementary modelling approaches have emerged: While so-called adaptive-walk models consider adaptation from the successive fixation of *de-novo* mutations only, quantitative genetic models assume that adaptation proceeds exclusively from pre-existing standing genetic variation. The latter approach, however, has focused on short-term evolution of population means and variances rather than on the statistical properties of adaptive substitutions. Our aim is to combine these two approaches by describing the ecological and genetic factors that determine the genetic basis of adaptation from standing genetic variation in terms of the effect-size distribution of individual alleles. Specifically, we consider the evolution of a quantitative trait to a gradually changing environment. By means of analytical approximations, we derive the distribution of adaptive substitutions from standing genetic variation, that is, the distribution of the phenotypic effects of those alleles from the standing variation that become fixed during adaptation. Our results are checked against individual-based simulations. We find that, compared to adaptation from *de-novo* mutations, (i) adaptation from standing variation proceeds by the fixation of more alleles of small effect; (ii) populations that adapt from standing genetic variation can traverse larger distances in phenotype space and, thus, have a higher potential for adaptation if the rate of environmental change is fast rather than slow.

## INTRODUCTION

One of the biggest surprises that has emerged from evolutionary research in the past few decades is that, in contrast to what has been claimed by the neutral theory (Kimura 1983), adaptive evolution at the molecular level is wide-spread. In fact, some empirical studies concluded that up to 45% of all amino acid changes between *Drosophila simulans* and *D. yakuba* are adaptive (Smith and Eyre-Walker 2002; Orr 2005b). Along the same line, Wichman *et al.* (1999) evolved the single-stranded DNA bacteriophage ΦX174 to high temperature and a novel host and found that 80 – 90% of the observed nucleotide substitutions had an adaptive effect. These and other results have led to an increased interest in providing a formal framework for the adaptive process that goes beyond traditional population- and quantitative-genetic approaches and considers the statistical properties of suites of substitutions in terms of “individual mutations that have individual effects” (Orr 2005a). In general, selection following a change in the environmental conditions may act either on *de-novo* mutations or on alleles already present in the population, also known as standing genetic variation. Consequently, from the numerous studies that have attempted to address this subject, two complementary modelling approaches have emerged.

So-called adaptive-walk models (Gillespie 1984; Kauffman and Levin 1987; Orr 2002, 2005b) typically assume that selection is strong compared to mutation, such that the population can be considered monomorphic all the time and all observed evolutionary change is the result of *de-novo* mutations. These models have produced several robust predictions (Orr 1998, 2000; Martin and Lenormand 2006a,b), which are supported by growing empirical evidence (Cooper *et al.* 2007; Rockman 2012; Hietpas *et al.* 2013; but see Bell 2009), and has provided a statistical framework for the fundamental event during adaptation, that is, the substitution of a resident allele by a beneficial mutation. Specifically, the majority of models (e.g., Gillespie 1984; Orr 1998; Martin and Lenormand 2006a) consider the effect-size distribution of adaptive substitutions following a sudden change in the environ-ment. Recently, Kopp and Hermisson (2009b) and Matuszewski *et al.* (2014) extended this framework to gradual environmental change.

In contrast, most quantitative-genetic models consider an essentially inexhaustible pool of pre-existing standing genetic variants as the sole source for adaptation (Lande 1976). Evolving traits are assumed to have a polygenic basis, where many loci contribute small individual effects, such that the distribution of trait values approximately follows a Gaussian distribution (Bulmer 1980; Barton and Turelli 1991; Kirkpatrick *et al.* 2002). Since the origins of quantitative genetics lie in the design of plant and animal breeding schemes (Wricke and Weber 1986; Tobin *et al.* 2006; Hallauer *et al.* 2010), the traditional focus of these models was on predicting short-term changes in the population mean phenotype (often assuming constant genetic variances and covariances), and not on the fate and effect of individual alleles. The same is true for the relatively small number of models that have studied the contribution of new mutations in the response to artificial selection (e.g., Hill and Rasbash 1986a) and the shape and stability of the G-matrix (i.e., the additive variance-covariance matrix of genotypes; Jones *et al.* 2004, 2012).

It is only in the past decade that population geneticists have thoroughly addressed adaptation from standing genetic variation at the level of individual substitutions (Orr and Betancourt 2001; Hermisson and Pennings 2005; Chevin and Hospital 2008). Hermisson and Pennings (2005) calculated the probability of adaptation from standing genetic variation following a sudden change in the selection regime. They found that, for small-effect alleles, the fixation probability is considerably increased relative to that from new mutations. Furthermore, Chevin and Hospital (2008) showed that the selective dynamics at a focal locus are substantially affected by genetic background variation. These results where experimentally confirmed by Lang *et al.* (2011), who followed beneficial mutations in hundreds of evolving yeast populations and showed that the selective advantage of a mutation plays only a limited role in determining its ultimate fate. Instead, fixation or loss is largely determined by variation in the genetic background – which need not to be preexisting, but could quickly be generated by a large number of new mutations. Still, predictions beyond these single-locus results have been verbal at best, stating that “compared with new mutations, adaptation from standing genetic variation is likely to lead to faster evolution [and] the fixation of more alleles of small effect […]” (Barrett and Schluter 2008). Thus, despite recent progress, one of the central questions still remains unanswered: From the multitude of standing genetic variants segregating in a population, which are the ones that ultimately become fixed and contribute to adaptation, and how does their distribution differ from that of (fixed) *de-novo* mutations?

The aim of the present article is to contribute to overcoming what has been referred to as “the most obvious limitation” (Orr 2005b) of adaptive-walk models and to study the ecological and genetic factors that determine the genetic basis of adaptation from standing genetic variation. Specifically, we consider the evolution of a quantitative trait in a gradually changing environment. We develop an analytical framework that accurately describes the distribution of adaptive substitutions from standing genetic variation and discuss its dependence on the effective population size, the strength of selection and the rate of environmental change. In line with Barrett and Schluter (2008), we find that, compared to adaptation from *de-novo* mutations, adaptation from standing genetic variation proceeds, on average, by the fixation of more alleles of small effect. Furthermore, when standing genetic variation is the sole source for adaptation, faster environmental change can enable the population to remain better adapted and to traverse larger distances in phenotype space.

## MODEL AND METHODS

### Phenotype, Selection and Mutation

We consider the evolution of a diploid population of *N* individuals with discrete and nonoverlapping generations characterized by a single phenotypic trait *z*, which is under Gaussian stabilizing selection with regard to a time-dependent optimum *z*_opt_(*t*):

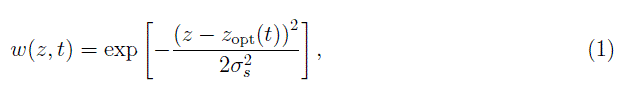

where 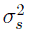 describes the width of the fitness landscape. Throughout this paper we choose the linearly moving optimum,

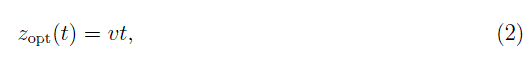

where *v* is the rate of environmental change.

Mutations enter the population at rate 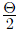 (with Θ = 4*Nu* where *u* is the per-haplotype mutation rate), and we assume that their phenotypic effect size *α* follows a Gaussian distribution with mean 0 and variance 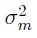 (which we will refer to as the distribution of new mutations), that is

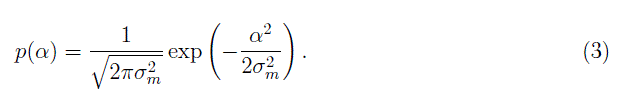

Throughout this paper we equate genotypic with phenotypic values and, thus, neglect any environmental variance. Note that this model is, so far, identical to the moving-optimum model proposed by Kopp and Hermisson (2009b) (see also Bürger 2000).

### Genetic assumptions and simulation model

To study the distribution of adaptive substitutions from standing genetic variation, we conducted individual-based simulations (IBS; available upon request; see Bürger 2000; Kopp and Hermisson 2009b) that explicitly model the simultaneous evolution at multiple loci, while making additional assumptions about the genetic architecture of the selected trait, the life cycle of individuals and the regulation of population size. This will serve as our main model.

**Genome** Individuals are characterized by a linear (continuous) genome of diploid loci, which determine the phenotype *z* additively (i.e., there is no phenotypic epistasis; note, however, that there is epistasis for fitness). Mutations occur at constant rate 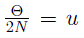 per haplotype. In contrast to the majority of individual-based models (e.g., Jones *et al.* 2004; Kopp and Hermisson (2009b); Matuszewski *et al.* 2014), we do not fix the number of loci *a-priori*, but instead assume that each mutation creates a unique polymorphic locus, whose position is drawn randomly from a uniform distribution over the entire genome (where genome length is determined by the recombination parameter *r* described below). Thus, each locus consists only of a wild-type allele with phenotypic effect 0 and a mutant allele with phenotypic effect α, which is drawn from equation (3). Thus, we effectively design a bi-allelic infinite-sites model with a continuum of alleles.

To monitor adaptive substitutions, we introduce a population-consensus genome 𝒢 that keeps track of all loci, that is, of all mutant alleles that are segregating in the population. If a mutant allele becomes fixed in the population it is declared the new wild-type allele and its phenotypic effect is reset to 0. The phenotypic effects of all fixed mutations are taken into account by a variable *z*_fix_, which can be interpreted as a phenotypic baseline effect. Thus, the phenotype *z* of an individual *i* is given by

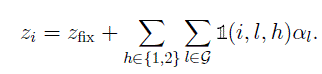

where

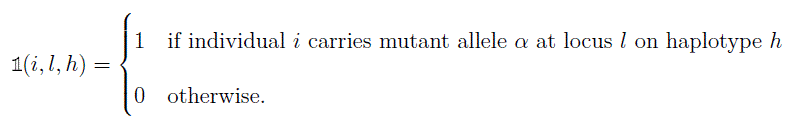

**Life cycle** Each generation, the following steps are performed:

1. *Viability selection:* Individuals are removed with probability 1 — *w(z)* (see (eq. 1).
2. *Population regulation:* If, after selection, the population size *N* exceeds the carrying capacity *K, N – K* randomly chosen individuals are removed.
3. *Reproduction:* The surviving individuals are randomly assigned to mating pairs, and each mating pair produces exactly *2B =* 4 offspring. Note that under this scheme, the effective population size *N_e_* equals 4/3 times the census size (Bürger 2000, p. 274). To account for this difference, Θ in the analytical approximations needs to calculated on the basis of this effective size, i.e., Θ = 4*N_e_u*. The offspring genotypes are derived from the parent genotypes by taking into account segregation, recombination and mutation.

**Recombination** For each reproducing individual, the number of crossing-over events during gamete formation (i.e., the number of recombination breakpoints) is drawn from a Poisson distribution with (genome-wide recombination) parameter *r* (i.e., the total genome length is *r* · 100cM, see Supporting Information 1). The genomic position of each recombination breakpoint is then drawn from a uniform distribution over the entire genome, and the offspring haplotype is created by alternating between the maternal and paternal haplotype depending on the recombination breakpoints. Free recombination (where all loci are assumed to be unlinked) corresponds to *r* → *∞*. In this case, for each locus a Bernoulli-distributed random number is drawn to determine whether the offspring haplotype will receive the maternal or the paternal allele at that locus.

**Simulation initialization and termination** Starting from a population of *K* wild-type individuals with phenotype *z* = 0 (i.e., the population was perfectly adapted at *t* = 0), we allowed for the establishment of genetic variation, 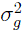, by letting the population evolve for 10,000 generations under stabilizing selection with a constant optimum. Increasing the number of generations had no effect on the average 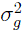. Following this equilibration time, the optimum started moving under ongoing mutational input, and the simulation was stopped once all alleles from the standing genetic variation had either been fixed or lost (i.e., when 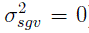. Simulations were replicated until a total number of 5000 adaptive substitutions from standing genetic variation was recorded.

### Analytical approximations: Evolution of a focal locus in the presence of genetic background variation

In order to obtain an analytically tractable model, we need to approximate the multi-locus dynamics. Clearly, simple interpolation of single locus theory will fail, because when alleles at different loci influencing the same trait segregate in the standing genetic variation, the selective dynamics of any individual allele are critically affected by the collective evolutionary response at other loci. In particular, any allele that brings the mean phenotype closer to the optimum simultaneously decreases the selective advantage of other such alleles (epistasis for fitness). Thus, if simultaneous evolution at many loci allows the population to closely follow the optimum, large-effect alleles at any given locus are likely to remain deleterious (as their carriers would overshoot the optimum). To account for these effects, we adopt a quantitative-genetics approach originally developed by Lande (1983) and introduce a genetic background *z*_B_ that evolves according to Lande’s equation

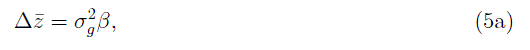

where

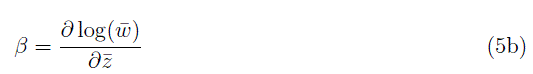

denotes the selection gradient, which measures the change in log mean fitness per unit change of the mean phenotype and 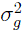 gives the genetic variance (Lande 1976). Furthermore, assuming that the distribution of phenotypic values from the genetic background is Gaussian and the genetic variance remains constant, the mean background phenotype evolves according to

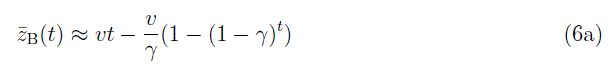

with

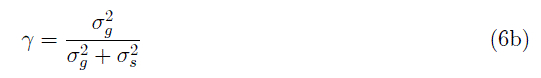

(Bürger and Lynch 1995).

Given the dynamics of the genetic background, we choose one focal locus and derive the time-dependent selection coefficient *s*(α, *t*) for an allele with phenotypic effect α (for details see below). We then use theory for adaptation from standing genetic variation (Hermisson and Pennings 2005) and for fixation under time-inhomogeneous selection (Uecker and Hermisson 2011) to estimate the fixation probability for this allele (see also Appendix 1). As long as there is no linkage (i.e., there is free recombination between *all* loci), each locus can be viewed as the focal locus (with a specific phenotypic effect α), allowing us to get an estimate for the overall distribution of adaptive substitutions from standing genetic variation. Thus, in these approximations, our multi-locus model is effectively treated within a single-locus framework. Note that a similar focal-locus approach has recently been used to analyze the effect of genetic background variation on the trajectory of an allele sweeping to fixation (Chevin and Hospital 2008), and to study the probability of adaptation to novel environments (Gomulkiewicz *et al.* 2010), with both studies stressing the fact that genetic background variation cannot be neglected and critically affects the adaptive outcome.

### Wright-Fisher simulations: A focal locus with recurrent mutations

To simulate evolution at a focal locus, we followed Hermisson and Pennings (2005) and implemented a multinomial Wright-Fisher (WF) sampling approach (available upon request). These simulations serve as an additional analysis tool that has been adjusted to the approximation method and allows the adaptive process to be simulated fast and efficiently. In addition, they go beyond the individual-based model in one aspect, as they do not make the infinite-sites assumption but allow for recurrent mutation at the focal locus.

**Genome** At the focal locus, mutations with a fixed allelic effect *α* appear recurrently at rate *θ* and convert ancestral alleles into derived mutant alleles. Accordingly, despite a genetic background with normally distributed genotypic values, there are at most two types of (focal) alleles in the population, where each type “feels” only the mean background *z̄*_B_, which evolves according to Lande’s equation (eq. 5, see above). The genetic background variation 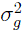 is assumed to be constant and serves as a free parameter that is independent of θ, *N*_*e*_ and 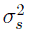. Note that the evolutionary response at the focal locus is influenced by that of the genetic background, and vice versa, meaning that the two are interdependent.

**Procedure** We follow the evolution of 2*N_e_* alleles at the focal locus. Each generation is generated by multinomial sampling, where the probability of choosing an allele of a given type (ancestral or derived) is weighted by its respective (marginal) fitness. Furthermore, the mean phenotype of the genetic background *z̄*_B_ evolves deterministically according to equation (5) with constant 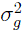. To let the population reach mutation-selection-drift equilibrium, each simulation is started 4*N_e_* generations before the environment starts changing. Initially, the population consists of only ancestral alleles “0”; the derived allele “1” is created by mutation

If the derived allele reaches fixation before the environmental change (by drift), it is itself declared “ancestral”; i.e., the population is set back to the initial state. After 4*N_e_* generations, the optimum starts moving, such that the selection coefficient of the derived allele, which is initially deleterious (i.e., *s*(*α, t*) ≤ 0), increases and may eventually become beneficial (i.e., *s*(*α, t* > 0), depending on the response at the genetic background. Simulations continue until the derived allele is either fixed or lost. Fixation probabilities are estimated from 100,000 simulation runs.

Both simulation programs are written in C++and make use of the Gnu Scientific Library (Galassi *et al.* 2009). Mathematica (Wolfram Research, Inc., Champaign, USA) was used for the numerical evaluation of integrals and to create plots and graphics, making use of the LevelScheme package (Caprio 2005).

A summary of our notation is given in Table 1.

**Table 1.** A summary of notation and definitions.

|  |  |
| --- | --- |
| $\alpha$ | phenotypic effect of mutation |
| $p(\alpha)$ | (Gaussian) distribution of new mutations |
| $z$ | phenotype |
| $\bar{z}_B$ | mean genetic background phenotype |
| $v$ | rate of environmental change |
| $w(z, z_{opt}(t))$ | (Gaussian) fitness function |
| $\sigma_s^2$ | width of Gaussian fitness function |
| $\sigma_m^2$ | variance of new mutations |
| $\sigma_g^2$ | (background) genetic variance |
| $s(\alpha, t)$ | time-dependent selection coefficient for allele with phenotypic effect $\alpha$ |
| $x$ | frequency of mutant allele |
| $N_e$ | effective population size |
| $\theta$ | per locus mutation rate |
| $\Theta$ | population-wide mutation rate (per trait) |
| $\Pi_{fix}$ | fixation probability |
| $\rho(x, \alpha)$ | Distribution of mutant allele frequency at a single locus with phenotypic effect $\alpha$ |
| $P_{SGV}$ | Probability to adapt from standing genetic variation |
| $p_{SGV}$ | Distribution of adaptive substitutions from standing genetic variation |
| $\delta_{eq}$ | equilibrium lag |

## RESULTS

In the following, we calculate, first, the probability that a focal allele from the standing genetic variation becomes fixed when the population adapts to a moving phenotypic optimum, and second, the effect-size distribution of such alleles. Note that the first result will be derived under the assumption of recurrent mutation (see “Wright-Fisher simulations”), and serves as an intermediate step for the second result, which is based on an infinite-sites model (see “Genetic assumptions and simulation model”).

### The probability for adaptation from standing genetic variation

The probability that a focal mutant allele from the standing genetic variation contributes to adaptation depends on the dynamics of its the selection coefficient in the presence of genetic background variation. For an allele with effect α and a genetic background with mean *z̄*_B_ and variance 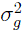, the selection coefficient can be calculated as

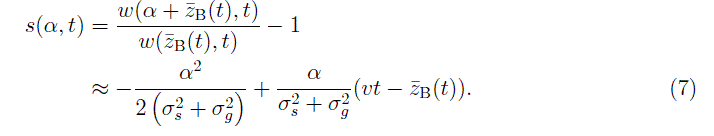

Note that the genetic background variance has the effect of broadening the fitness landscape experienced by the focal allele (the term 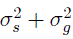.

Plugging equation (6a) into equation (7) then yields the selection coefficient,

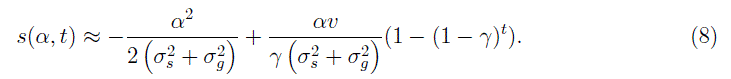

Assuming that the population is perfectly adapted at *t* = 0 (*z̄_B_* = 0), the initial (deleterious) selection coefficient is given by

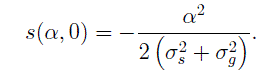

Unlike in the model without genetic background variation (Kopp and Hermisson (2009b)), *s(α, t)* does not increase linearly, but instead depends on the evolution of the phenotypic lag *δ* between the optimum and the mean background phenotype. In particular, the population will reach a dynamic equilibrium with Δ*z̄*_B_ = *v*, where it follows the optimum with a constant lag

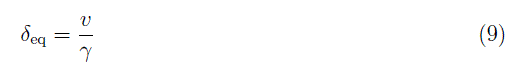

(Bürger and Lynch 1995). Consequently, the selection coefficient for α approaches

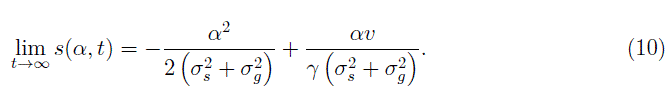

Note that the right-hand side can be written as *s*(*α*, 0) + α*β*_eq_, where *β*_eq_ is the equilibrium selection gradient (Kopp and Matuszewski 2014). In this case, the largest obtainable selection coefficient is for α = δ_eq_ and evaluates to

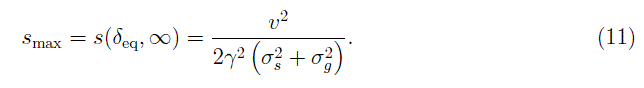

The range of allelic effects α that can reach a positive selection coefficient is bounded by α_min_ = 0 and α_max_ = 2δ_eq_. Note that in previous adaptive-walk models (e.q., Kopp and Hermisson (2009b); Matuszewski *et al.* 2014) there was no strict α_max_, since the population followed the optimum by stochastic jumps, whereas in the present model, the genetic background evolves deterministically and establishes a constant equilibrium lag.

Assuming that α was deleterious prior to the environmental change, its allele frequency spectrum *ρ*(*x*, *α*) is given by equation (A5). When genetic background variation is absent the fixation probability Π_fiz_(α) (eq. A7) can be calculated explicitly using

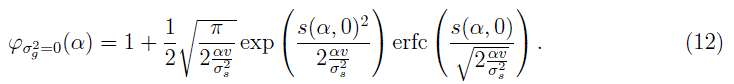

For the general case, however, Π_fix_(α) can only be calculated numerically using equation (8) in equation (A7b), yielding

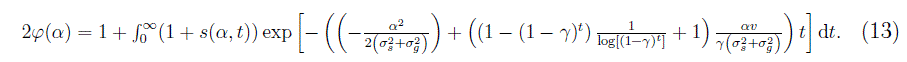

The fixation probability for an allele from the standing genetic variation with allelic effect α and a recurrent (per locus) mutation rate *θ* can then be calculated as

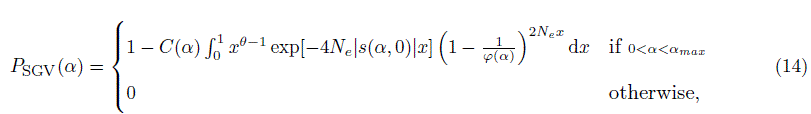

where 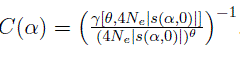.

When checked against Wright-Fisher simulations (see Methods for details), our analytical approximation equation (14) performs generally very well (Figs. 1 and S3_1). The only exception occurs when the background variation is high (large 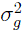) and stabilizing selection is weak (i.e., if 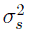 is large). In this case, equation (14) underestimates *P*_SGV_(α) for small α ∼ 0.5σ_*m*_. The reason is that, under a constant optimum (i.e., before the environmental change), the genetic background compensates for the deleterious effect of α (i.e., *z̄*_*B*_ < 0, in violation of our assumption that z̄_B_(0) = 0), effectively reducing the selection strength against the deleterious mutant allele. Consequently, α is, on average, present at higher initial frequencies than predicted by equation (A5).

Note that, if α is small compared to the genetic background variation (i.e., in the limit of α / σ_*m*_ → 0) and environmental change is slow (i.e., *v* ≪ 10^−5^), *P*_SGV_(α) will approach the probability of fixation from standing genetic variation for a neutral allele (i.e., α = 0), which can be calculated as

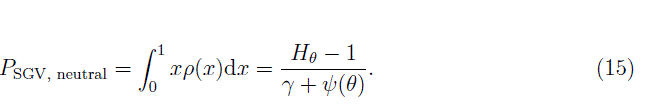

where *ρ(x)* is given by equation (A3), *H*_*n*_ denotes the *n*^th^ harmonic number, *γ* ≈ 0.577 is Euler’s gamma and *ψ(·)* is the polygamma function (see dashed lines in Figs. 1 and S3_1).

**Figure 1.**
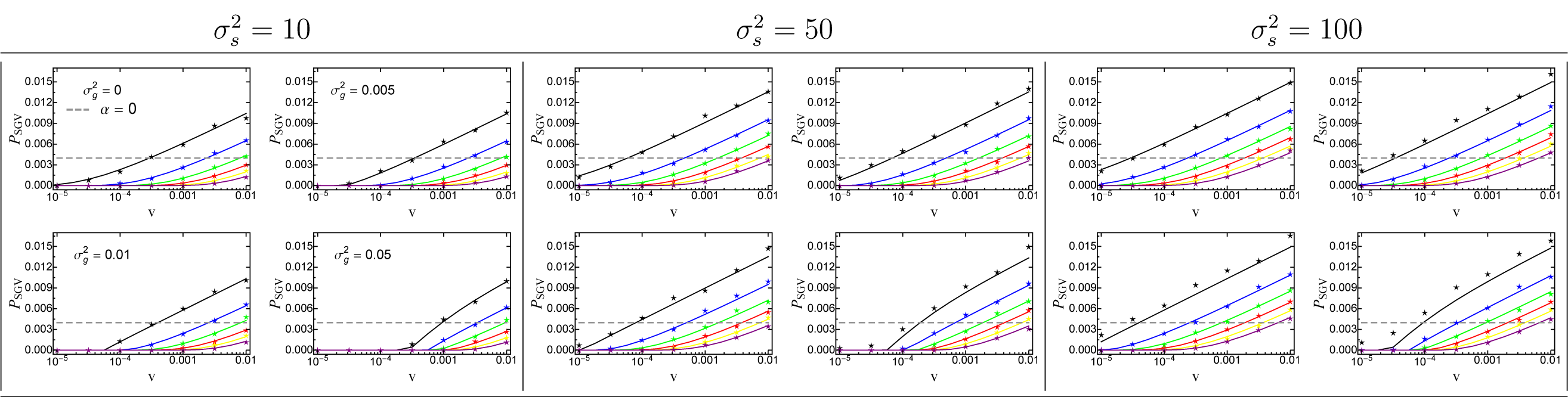
The probability for a mutant allele to adapt from standing genetic variation as a function of the rate of environmental change *v*. Solid lines correspond to the analytical prediction (eq. 14), the grey dashed line shows the probability for a neutral allele (*α* = 0; eq. 15), and symbols give results from Wright-Fisher simulations. The phenotypic effect size α of the mutant allele ranges from 0.5σ_m_ (top line; black) to 3σ_m_ (bottom line; purple) with increments of 0.5σ_m_. The figures in each parameter box (per locus mutation rate *θ*, width of fitness landscape 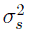 correspond to different values of the genetic background variation 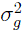 with 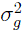 = 0 (no background variation; top left), 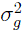 *=* 0.005 (top right), 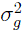 *=* 0.01 (bottom left) and 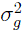 = 0.05 (bottom right). Other parameters: *N*_*e*_ = 25000, *θ* = 0.004, 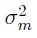 =0.05.

Figures 1 and S3_1 show some general trends: First, the probability for a mutant allele tobecome fixed increases with the rate of environmental change, *v*, (irrespective of its effect size α, the per locus mutation rate *θ* and the width of the fitness landscape 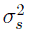) since only the positive term in equation (8) depends (linearly) on *v*. Second, *P*_SGV_(α) is proportionalto *θ* as long as *θ* is small (compare *θ =* 0.004 and *θ =* 0.04 in Fig. S3_1), simply because theprobability that α is present in the population is linear in *θ*. Thus, Figure 1 is representative for the limit *θ →* 0 which will be used below. Indeed, only if the per-locus mutation rate is fairly large (*θ* > 0.1), does the shape of the distribution of allele frequencies becomeimportant, and the increase in *P*_SGV_(α) with *θ* becomes less than linear (Fig. S3_1). Third,changes in the width of the fitness landscape, 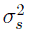, have a dual effect: While increasing 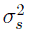 promotes the initial frequency of the focal allele in the standing genetic variation (because stabilizing selection is weaker), the selection coefficient increases more slowly after the onset of environmental change (such that the allele is less likely to be picked up by selection; see eq. 7). Our results, however, show that the former effect always outweighs the latter (as *P*_SGV_(α) increases with 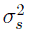). Finally, if the genetic background variation 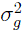 is below a threshold value (e.g., 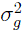 < 0.005; the exact threshold should depend on *θ* and 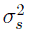) it only marginally affects the fixation probability of the focal allele α. Once 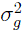 surpasses this value, however, it critically affects *P*_SGV_(α) (in accordance with the results by Chevin and Hospital 2008). In particular, as 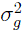 increases *P*_SGV_(α) decreases, because most large-effect alleles remain deleterious even if environmental change is fast. Thus, enlarged background variation acts as if reducing the rate of environmental change *v*. In summary, our analytical results are in good agreement with the WF-simulation model, and will serve as an important first step towards deriving the distribution of adaptive substitutions from standing genetic variation.

### The distribution of adaptive substitutions from standing genetic variation

We now derive the distribution of adaptive substitutions from standing genetic variation over all mutant effects α. In the previous section, we derived the fixation probability at a focal locus (with a given effect α) by treating the genetic background variance 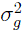 as an independent model parameter. In the full model, this variance results from a balance of mutation, selection and drift at all background loci. As such, it is a function of the basic model parameters for these forces. Since we use an infinite-sites model, there is no recurrent mutation and each allele originates from a single mutation. Consequently, the amount of background variation 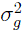 is accurately predicted by the Stochastic-House-of-Cards (SHC) approximation (not shown; Bürger and Lynch 1995)

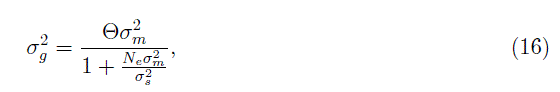

where mutation is parametrized by the total (per trait) mutation rate Θ and the mutational variance 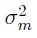, the width of the fitness landscape is given by 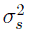, and the effective population size *N*_*e*_ is a measure for genetic drift.

To derive the probability that an allele with a given phenotypic effect α contributes to adaptation, we first need to calculate the probability that such an allele segregates in the population at time 0. Following Hermisson and Pennings (2005), the probability *P*_0_ that the allele is *not* present can be approximated by integrating over the distribution of allele frequencies *ρ(x*, α) (eq. A5) from 0 to 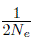 yielding

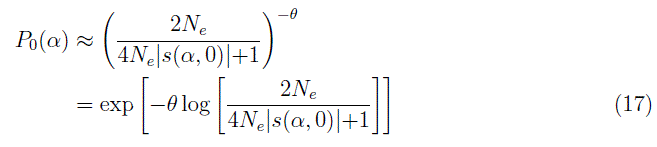

(eq. 7 and Appendix of Hermisson and Pennings 2005). The fixation probability can then be calculated by conditioning on segregation of the allele in the limit θ → 0 (due to the infinite-sites assumption). Using equation (14), this probability reads

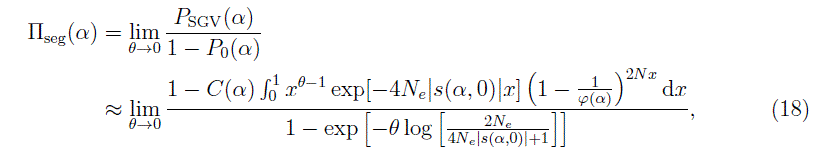

where 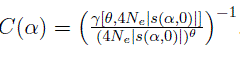 (see also eq. A5) and with *φ*(α) according to equation (13).

The limit in equation (18) can be approximated numerically by setting *θ* to a very small, but positive value.

Multiplying by the rate of mutations with effect α (i.e., Θρ(α)), the distribution of adaptive substitutions from standing genetic variation is given by

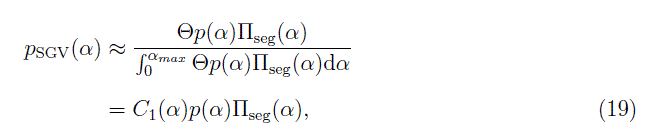

where C_1_(α) is a normalization constant (black line in Figs. 2, 3 and Fig. 4). Note that equation (19) still depends on Θ through its effect on the background variance 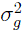 (which affects 𝚷_seg_(α)). In particular, in the SHC approximation (eq. 16), 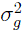 scales linearly with Θ. Furthermore, equation (19) should be valid for any distribution of mutational effects *p(α).*

In the limit where the equilibrium lag is reached fast (i.e., when γ is large; eq. 6b), the moving-optimum model reduces to a model with constant selection for any focal allele (i.e., as in Hermisson and Pennings 2005). Using equations (A6) and (17) the fixation probabilityfor a segregating allele can be calculated as

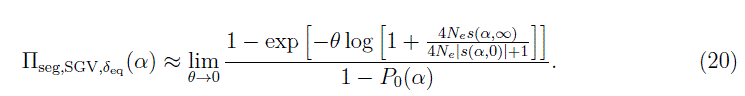

Plugging equation (20) into equation (19), the distribution of adaptive substitutions from standing genetic variation can be approximated by

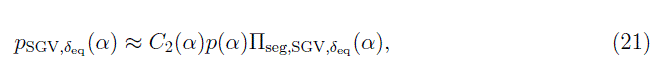

where *C*_*2*_(α) is a normalization constant (red line in Figs. 2, 3).

Similarly, the fixation probability of *de-novo* mutations under the equilibrium lag δ_eq_ can be derived (using 11 and eq. A2 with an initial frequency of 1/(2*N*)) as

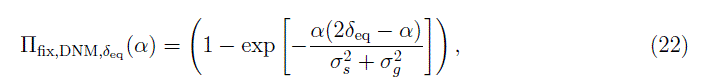

yielding the distribution of adaptive substitutions

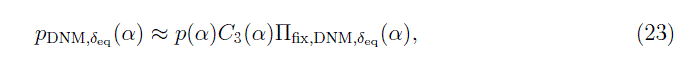

where *C*_*3*_(α) is a normalization constant (grey curve in Figs. 2, 3).

In contrast, if the environment changes very slowly, we can calculate the limit distribution of adaptive substitutions from standing genetic variation by approximating the fixation probability by that of a neutral allele (i.e., its allele frequency *x*). In this case,

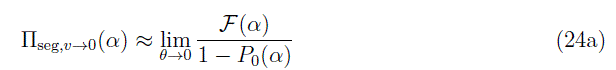

with

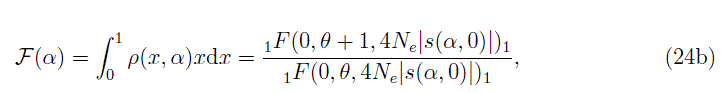

where *ρ(x, α)* is given by equation (A4) and the right-hand side is a ratio of hypergeometric functions.

Using equation (24a) the distribution of substitutions from standing genetic variation reads

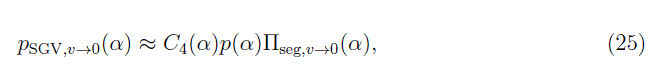

where *C*_4_(α) again denotes a normalization constant (blue line in Figs. 3, S3_2).

**The accuracy of the approximation** When compared to individual-based simulations, our analytical approximation for the distribution of adaptive substitutions from standing genetic variation (eq. 19) performs, in general, very well as long as selection is strong, that is, the rate of environmental change *v* is high and/or the width of the fitness landscape 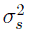 is not too large (Fig. 2). Under weak selection, however, equation (19) fails to capture the fixation of alleles with neutral or negative effects (“backward fixations”; *α* ≤ 0). The reason is that equation (A7) only considers the fixation of alleles whose selection coefficient *s(α,t)* becomes positive in the long term. But if the rate of environmental change is slow (or 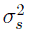 is very large), most alleles get fixed or lost simply by chance, that is, genetic drift. In particular, if genetic drift is the main driver of phenotypic evolution (i.e., *N_e_|s(α,t)| <* 1), the distribution of adaptive substitutions is almost symmetric around 0 (see Fig. S3_2). This distribution is described very well by equation (25), which assumes that the fixation probability of an allele is proportional to its initial frequency in the standing variation. In addition, even for cases where environmental change imposes modest directional selection, equation (25) still captures the shape of the distribution of adaptive substitutions reasonably well, when centered around the empirical mean (blue line in Figs. 2, 3).

With a moving phenotypic optimum, the selection coefficient (eq. 8) is initially very small. Accordingly, there is always a phase during the adaptive process where genetic drift dominates, that is, where *N_e_|s*(α,*t*)| < 1 for all mutant alleles. The length of this phase (i.e., the time it takes until selection becomes the main force of evolution) depends on the interplay of multiple parameters, notably *v* 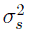, *N*_*e*_ and Θ. A good heuristic to determine whether evolution will ultimately become dominated by selection is to calculate *N*_*e*_*s*_max_ (eq. 11), which gives the maximal population-scaled selection coefficient. Since the selection coefficient of most mutations will be smaller than this value, one can consider as a rule of thumb that selection is the main driver of evolution as long as *N*_*e*_*s*_max_ ≥ 10. In this case, equation (19) matches the individual-based simulations very well (see asterisks in Figs. 2, 3). In summary, the accuracy of our approximation crucially depends on the efficacy of selection.

The effects of linkage on the distribution of adaptive substitutions from standing genetic variation are discussed in Supporting Information 1. The main result is that only tight linkage has a noticeable effect, namely to reduce the efficacy of selection and increase the proportion of “backward” fixations (moving the distribution closer to the prediction from eq. 25).

**Biological interpretation** As shown in Figures (2) and (3), adaptive substitutions from standing genetic variation have, on average, smaller phenotypic effects than those from *de-novo* mutations. There are two reasons for this result. First, in the standing genetic variation, small-effect alleles are more frequent than large-effect alleles and might already segregate at appreciable frequency (increasing their fixation probability). Second, substitutions from standing variation occur in the initial phase of the adaptive process, where the phenotypic lag is small, whereas our approximation for *de-novo* mutations (eq. 23) assumes that the phenotypic lag has reached its maximal (equilibrium) value (which need not be large, depending on the amount of genetic background variation). The relative importance of these two effects can be seen in Figures (2) and (3): Comparing the grey shaded area (eq. 23; *de-novo* mutations under the equilibrium lag) with the red line (eq. 21; standing genetic variation under the equilibrium lag) shows the effect of larger starting frequencies of small-effect mutations from the standing genetic variation. The difference of the black (eq. 19; standing genetic variation) and red (eq. 21; standing genetic variation under the equilibrium lag) lines show the effects of the initially smaller lag (i.e., the effect of the dynamical selection coefficient). Note that the first effect is always important (even if Θ and 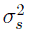 are large and *v* is small, where the red line and the grey curve almost coincide—though this is only because the approximation is bad). The second effect, however, becomes particularly important if 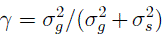 is small (i.e, if the time to reach the equilibrium lag is large), such that selection coefficients are dynamic and small-effect alleles are selected earlier than large-effect alleles, explaining the relative lack of large-effect alleles in the distribution of adaptive substitutions.

**Figure 2.**
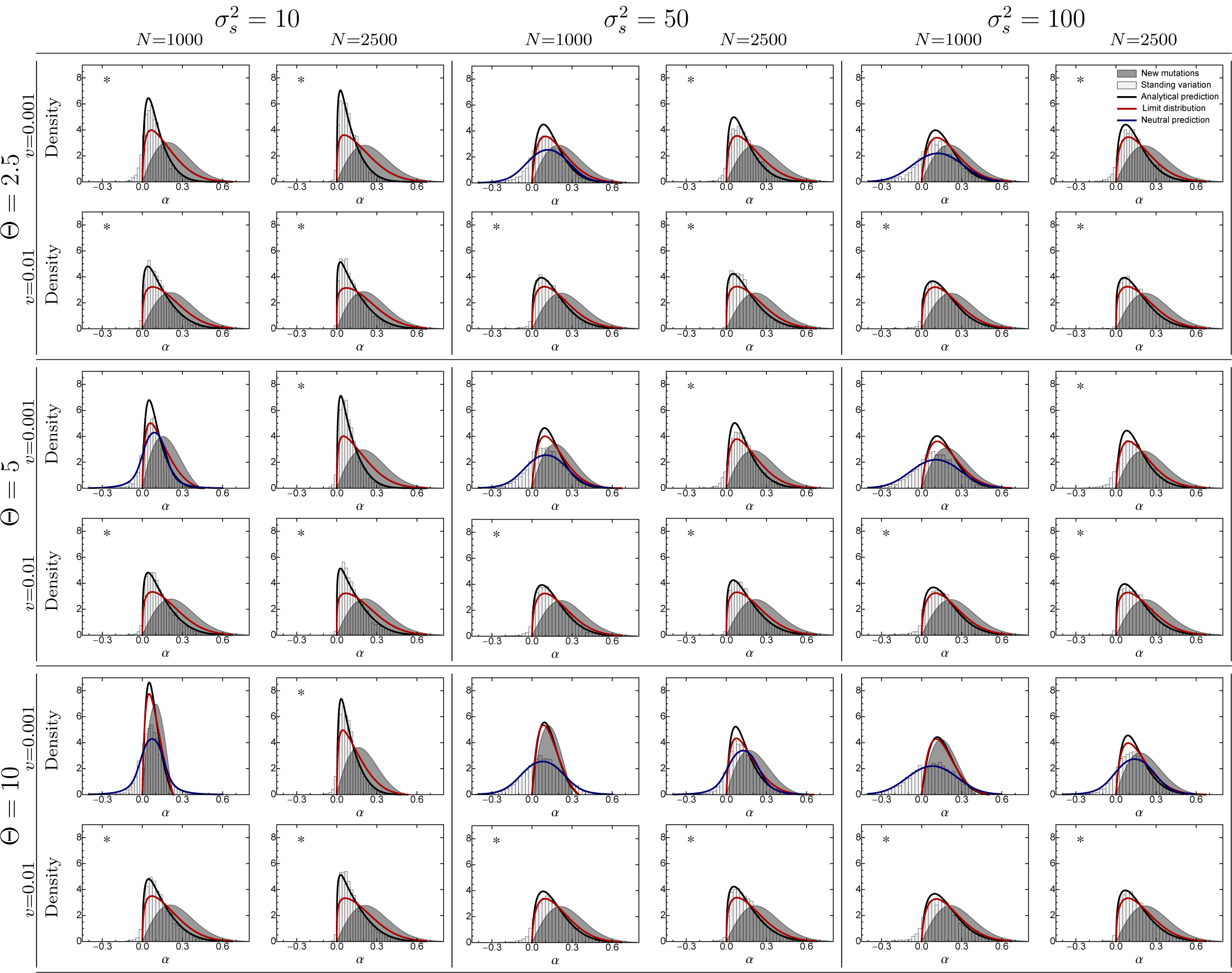
The distribution of adaptive substitutions from standing genetic variation. Histograms show results from individual-based simulations. The black line corresponds to the analytical prediction (eq. 19), with the genetic background variation 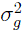 determined by the SHC approximation (eq. 16). The red line gives the analytical prediction for the limiting case where the equilibrium lag δ_eq_ is reached fast (eq. 21). The blue line is based on the analytical prediction Eq. (25) — which assumes a neutral fixation probability — but has been shifted so that it is centered around the empirical mean. The grey curve gives the analytical prediction for substitutions from *de-novo 396* mutations under the assumption that the phenotypic lag δ_eq_ has reached its equilibrium (eq. 23). The asterisks indicate where *N*_e_s_max_ ≥ 10. Fixed parameter: 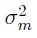=0.05.

**Figure 3.**
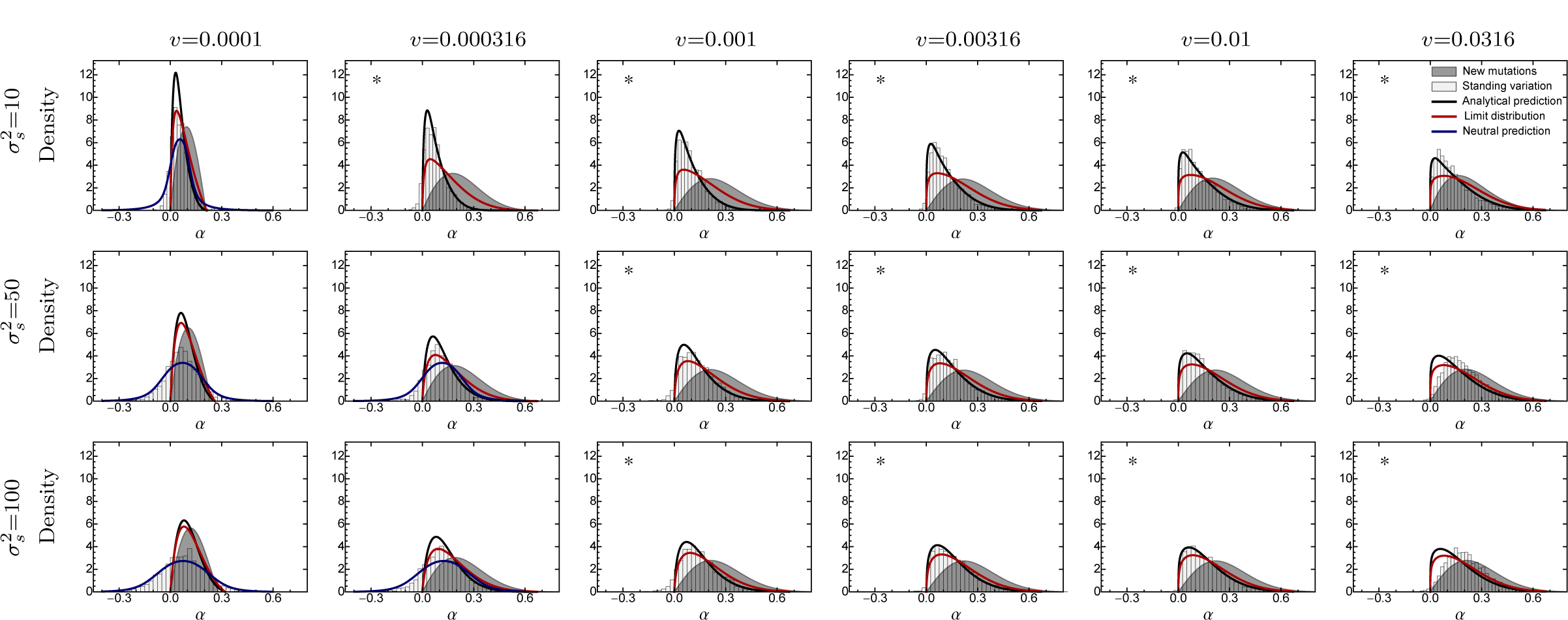
The distribution of adaptive substitutions from standing genetic variation for various rates of environmental change. For further details see Fig. 2. Fixed parameters: Θ = 2.5, *N* = 2500, 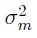 = 0.05.

Generally, the distribution of adaptive substitutions is unimodal and generally resembles a log-normal distribution (Figs. 2, 3). Only if selection is very weak (i.e., when 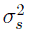 is large and/or *v* is small), does it contain a significant proportion of “backward fixations” (with negative *α*; Fig. 3; see “Accuracy of the Approximation”). As the rate of environmental change *v* increases, the mean phenotypic effect of substitutions increases (Fig. 4, top row), too, but the mode may actually decrease (Fig. 3), that is, the distribution becomes more asymmetric and skewed, resembling the “almost exponential” distribution of substitutions from *de-novo* mutations in the sudden change scenario (Orr 1998). A likely explanation is that small-effect alleles, which are common in the standing variation, are under stronger selection and have an increased fixation probability if *v* is large (see Fig. 1).

Interestingly, if the environment changes very fast the simulated distribution of adaptive substitutions from standing genetic variation almost exactly matches the one predicted by equation (23) for *de-novo* mutations (Fig. 5, see also Figs. 2, 3). However, this seems to be an artefact rather than a relevant biological phenomenon. The reason is that the environment changes so fast that the population quickly dies out. Thus, the resulting distribution of adaptive substitutions is that for a dying population and might not necessarily reflect the adaptive process. In an experimental setup, though, where populations evolve until they go extinct, the distribution of adaptive substitutions from standing genetic variation might truly be indistinguishable from that from *de-novo* mutations.

In the following, we discuss the influence of the other model parameters (Θ, 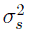 and *N*) on the distribution of adaptive substitutions from standing genetic variation, and in particular,its mean ᾱ (Fig. 4).

**Figure 4.**
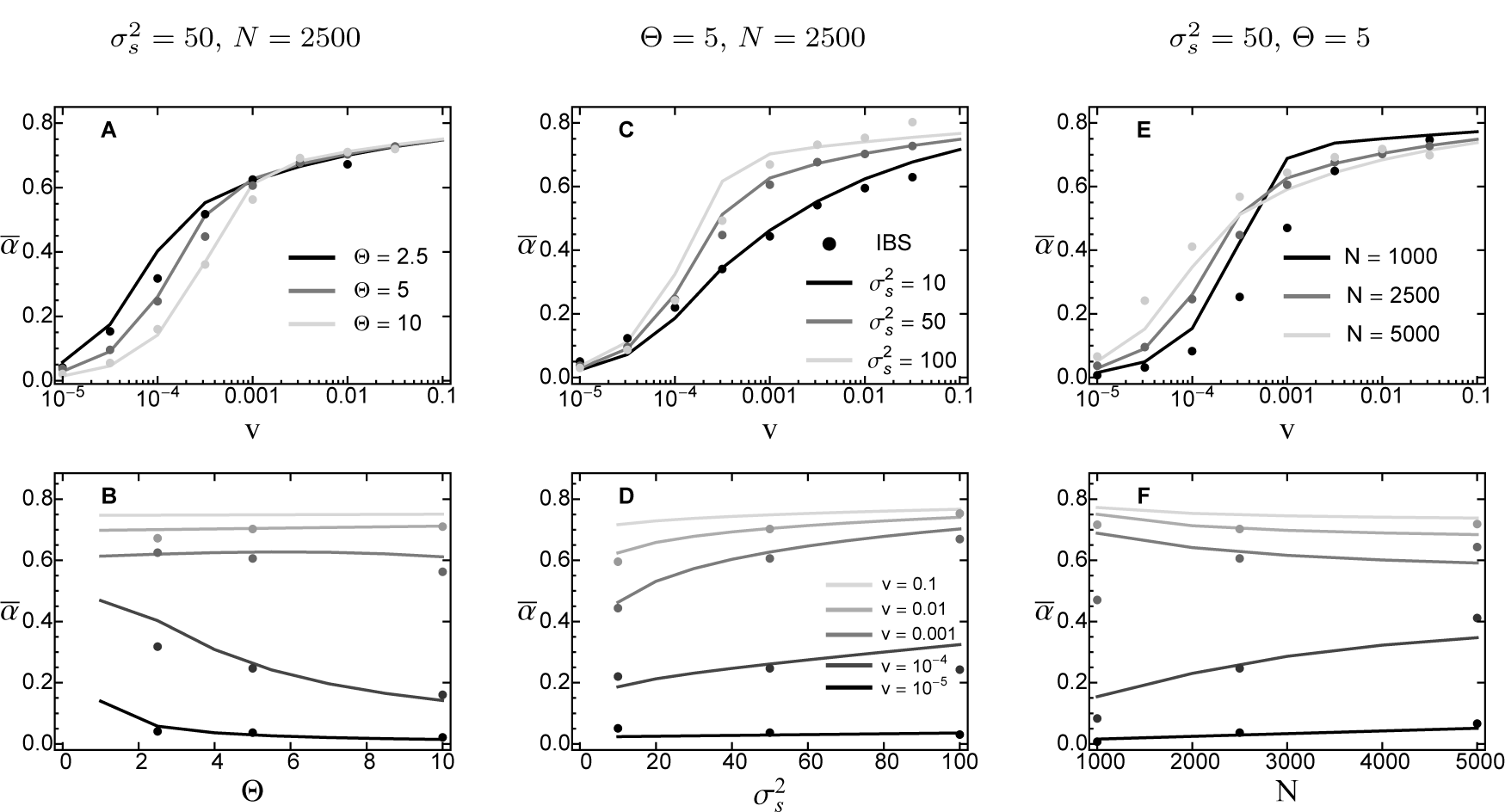
The mean size of adaptive substitutions from standing genetic variation, measured in units of mutational standard deviations (σ_m_) as a function of the rate of environmental change *v* (top row) and for various *v* as a function of the population-wide mutation rate Θ (bottom left), the width of the fitness landscape 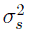 (bottom middle) and the population size *N* (bottom right). Lines show the analytical prediction (the mean of the distribution eq. eq:pDistMoveOpt), and symbols give results from individual-based simulations. Error bars for standard errors are contained within the symbols. For *v* = 0.1, no simulation results are shown, as these constitute a degenerate case (for details see “The accuracy of the approximation”). Fixed parameter: 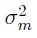 = 0.05.

**Figure 5.**
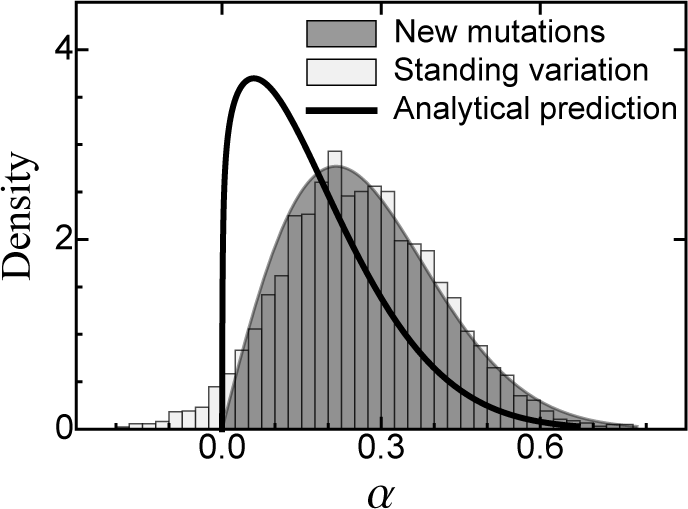
The distribution of adaptive substitutions from standing genetic variation in the case of fast environmental change. For further details see Fig. 2. Fixed parameters: 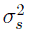 = 100, Θ = 10, *N = 2500, *v* = 0.1*, 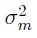 = 0.05.

The effect of the rate of mutational supply Θ depends strongly on the rate of environmental change *v: ᾱ* decreases with Θ if *v* is small but is independent of Θ if *v* is large (Fig. 4B). Recall that Θ enters *p*_SGV_(α) (eq. 19) only indirectly through the background variance 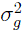. Accordingly, as Θ increases, so does 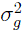 and, thus, *γ* (eq. 6b). In the limit *t →* ∞, the population will follow the optimum at a constant lag 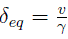. Thus, if *v* (such that, even for large 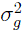, the lag is large relative to the mutational standard deviation σ_*m*_) increasing Θ does not affect ᾱ. In contrast, if *v* is small, increasing Θ (and, hence, 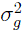 will reduce the lag even further, such that most large-effect alleles will be deleterious. Consequently, for small *v*, ᾱ decreases as Θ increases.

The width of the fitness landscape 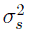 affects different aspects of the adaptive process, but its net effect is an increase of the mean effect size of fixed alleles as 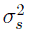 increases (i.e., as stabilizing selection gets weaker), especially if the rate of environmental change is intermediate (Fig. 4D). The reason is that weak stabilizing selection increases the frequency of large-effect alleles in the standing variation. In addition, weak selection also increases the phenotypic lag (eq. 9; see also Kopp and Matuszewski 2014), again favoring large effect alleles. Note that the latter point holds true even though weak selection increases the background variance 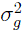. Finally, the effect of 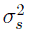 is strongest for intermediate *v*, because for small *v*, large-effect alleles are never favored, whereas for large *v*, all alleles with positive effect have a high fixation probability.

Similar arguments hold for *N*_*e*_ (when the rate of mutational supply, Θ, is held constant). First, increasing *N*_*e*_ will always increase the efficacy of selection, resulting in lower initial frequencies of mutant alleles (eq. A4) and decreased 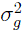 (eq. 16). If the environment changes slowly, ᾱ increases with *N*_*e*_, because the equilibrium lag increases (caused by the decrease in 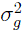. In contrast, if the rate of environmental change is fast, ᾱ slightly decreases with *N*_*e*_ due to the lower starting frequency of large-effect alleles and because small-effect alleles are selected more efficiently (i.e., they are less prone to get lost by genetic drift; Fig. 4F).

### The potential for adaptation from standing genetic variation and the rate of environmental change

So far, we have focussed on the distribution of adaptive substitutions for individual fixation events. We now address what can be said about the total progress that can be made from standing genetic variation following a moving phenotypic optimum. The overall potential for adaptation from standing genetic variation depends on the mean number of alleles segregating in the standing genetic variation, which can be accurately approximated as (Foley 1992)

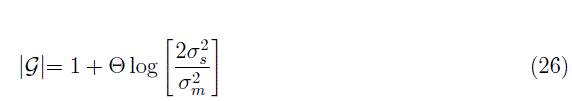

(results not shown). The mean number of alleles that become fixed can then be calculated as

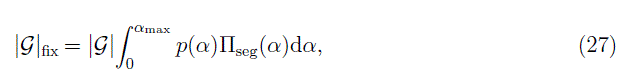

where the integral equals the normalization constant in equation (19) (i.e., the proportion of fixed alleles). Finally, using equation (27), the average distance travelled in phenotype space before standing variation is exhausted is given by

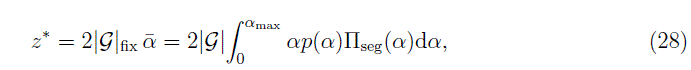

where ᾱ is the mean phenotypic effect size of adaptive substitutions from standing genetic variation, and the factor 2 in equation (28) comes from the fact that we are considering diploids (and α denotes the phenotypic effect per haplotype). Note that, once the shift of the optimum considerably exceeds *z*\*, the population will inevitably go extinct without the input of new mutations.

Figure 6 (see also Figs. S3_3, S3_4, S3_5 and Figs. S3_6, S3_7) illustrate these predictions and compare them to results from individual-base simulations (where, unlike in the rest of this paper, new mutational input was turned off after the onset of the environmental change).

**Figure 6.**
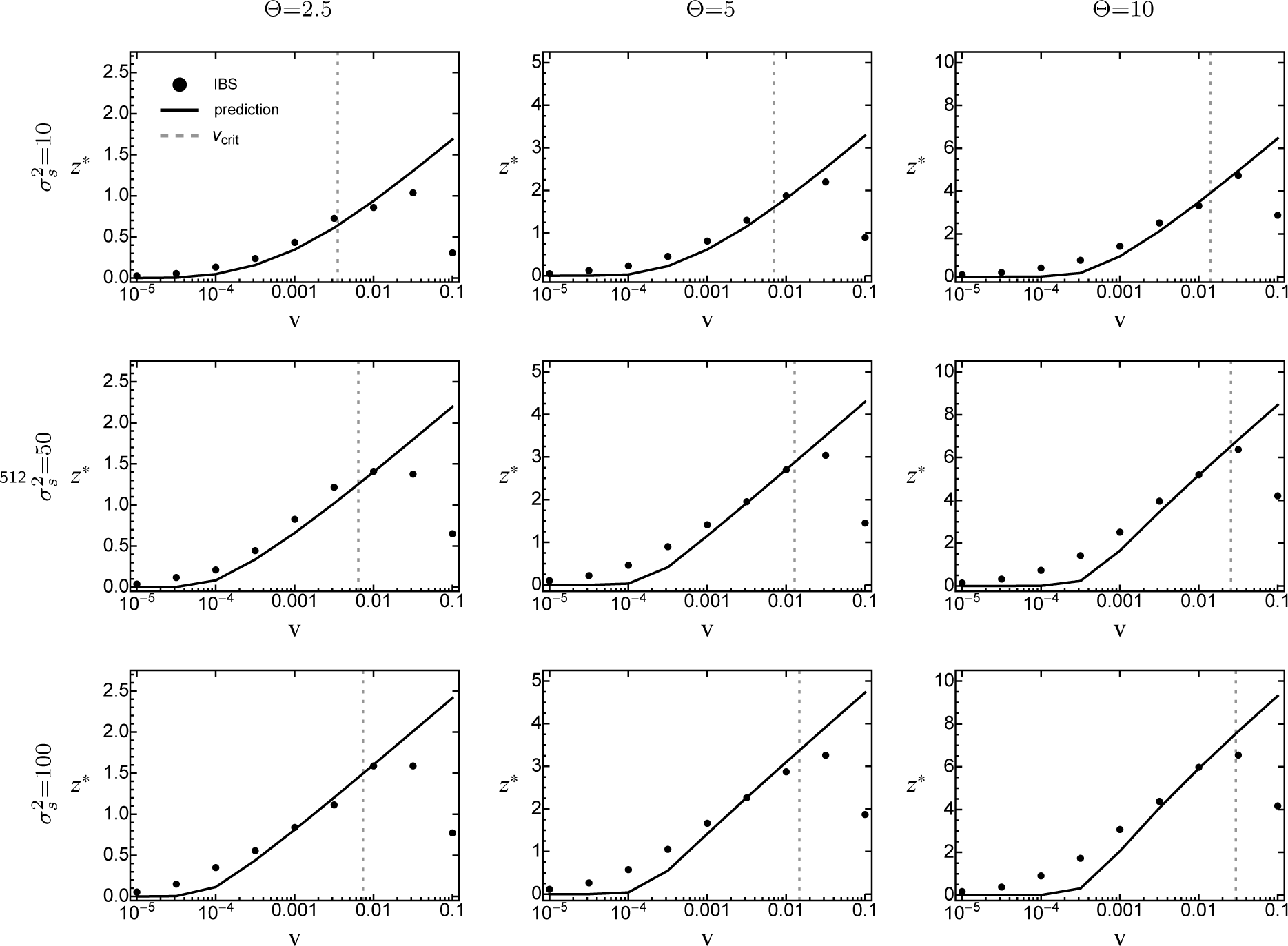
The average distance traversed in phenotype space, *z*^*^, as a function of the rate of environmental change *v*, when standing genetic variation is the sole source for adaptation. Symbols show results from individual-based simulations (averaged over 100 replicate runs). The black line gives the analytical prediction (eq. 28), with 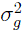 taken from equation (16). The grey-dashed line gives the critical rate of environmental change (eq. 29). Error bars for standard errors are contained within the symbols. Fixed parameters: *N* = 2500, 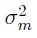 = 0.05.

Both the mean number of fixations |𝒢|_fix_ and the mean phenotypic distance travelled *z*\* increase with the rate of environmental change, reflecting the fact that more and larger-effect alleles become fixed if the environment changes fast. Only for very large *v*, where the rate of environmental change exceeds the “maximal sustainable rate of environmental change” (Bürger and Lynch 1995), which for our choice for the number of offspring *B* = 2 equals

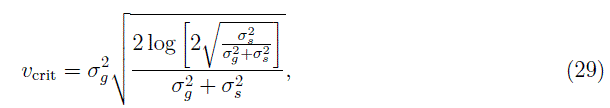

do |𝒢|_fix_ and *z*\* decrease sharply, because the population goes extinct before fixations can be completed (grey-dashed line in Figs. 6, S3_6 and S3_7). At small values of *v*, |𝒢|_fix_ matches the “neutral” prediction (grey-dashed line in Figs. S3_3, S3_4 and S3_5). Note that these fixations have almost no effect on *z*\*, because their average effect is zero. At intermediate *v*, equation (28) slightly underestimates *z*\* for parameter values leading to large background variance 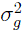 (i.e., high Θ and 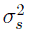). The likely reason is that the analytical approximation assumes 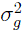 to be constant, while it obviously decreases in the simulations (since there are no *de-novo* mutations). All these results are qualitatively consistent across different values of 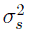 and Θ (Figs. 6, S3_6, S3_7).

### The relative importance of standing genetic variation and *de-novo* mutations over the course of adaptation

Until now we have compared adaptation from standing genetic variation to that from *de-novo* mutations in terms of their distribution of fixed phenotypic effects. We now turn to investigating their relative importance over the course of adaptation. For this purpose, we recorded (in individual-based simulations) the contributions of both sources of variation to the phenotypic mean and variance. An average time series for both measures is shown in Figure 7. As expected, the initial response to selection is almost entirely based on standing variation, but the contribution of *de-novo* mutations increases over time. As a quantitativemeasure for this transition, we define *t*_DNM,50_ (*z̄*) as the point in time where the cumulative contribution of *de-novo* mutations has reached 50%. Indeed, we find that, beyond this time, adaptation almost exclusively proceeds by the fixation of *de-novo* mutations (Fig. 7A). As expected, *t*_DNM,50_ (*z̄*) decreases with *v* (Figs. 8, S3_8, first row), while the total phenotypic response *z̄* increases (Figs. 8, S3_8, second row). The reason is that faster environmental change induces stronger directional selection and increases the phenotypic lag, such that standing variation is depleted more quickly and *de-novo* mutations and contribute earlier. Note that, as in Figure 6, the total phenotypic response at time *t*_DNM,50_ (*z̄*) decreases once *v* exceeds the “maximal sustainable rate of environmental change”, for the same reasons as discussed above. Furthermore, *t*_DNM,50_ (*z̄*) increases with both Θ and 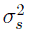 (due to the increased standing variation; see eq. 16). Interestingly, the relative contribution of original standing genetic variation to the total genetic variance at time *t*_DNM,50_ (*z̄*) remains largely constant (at around 20%) over large range of *v* and does not show any dependence on Θ nor 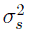 (Figs. 8, S3_8; third row). Deviations occur only if *v* is either very small or very large. In particular, if *v* is small, standing variation is almost completely depleted before new mutations play a significant role. Conversely, if *v* is very large, standing genetic variation still forms the majority of the total genetic variance. As mentioned above, this is most likely because the population goes extinct before fixations can be completed, that is, before the entire (standing) adaptive potential is exhausted. All these results remain qualitatively unchanged if, instead of t_DNM,50_ (*z̄)*, we define *t*_DNM,50_ (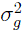) as the point in time where 50% of the current genetic variance goes back to *de-novo* mutations (Figs. S3_9, S3_10).

**Figure 7.**
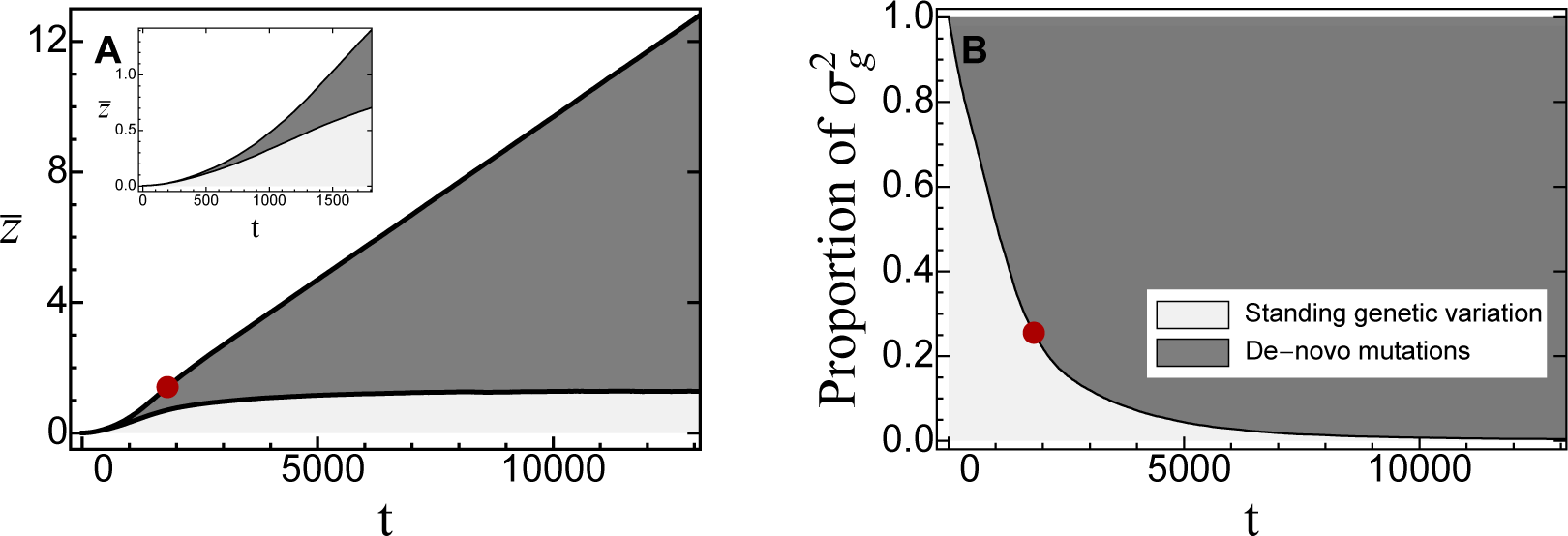
The contributions of standing genetic variation (light grey) and *de-novo* mutations (dark grey) to the cumulative phenotypic response to selection *z* (A) and the *current* genetic variance (B) over time. Plots show average trajectories over 1000 replicate simulations. The red dot marks the point in time where 50% of the total phenotypic response were due to *de-novo* mutations. The inset in (α) shows a more detailed plot of the dynamics of *z̄* up to this point. Fixed parameters: 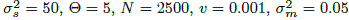.

**Figure 8.**
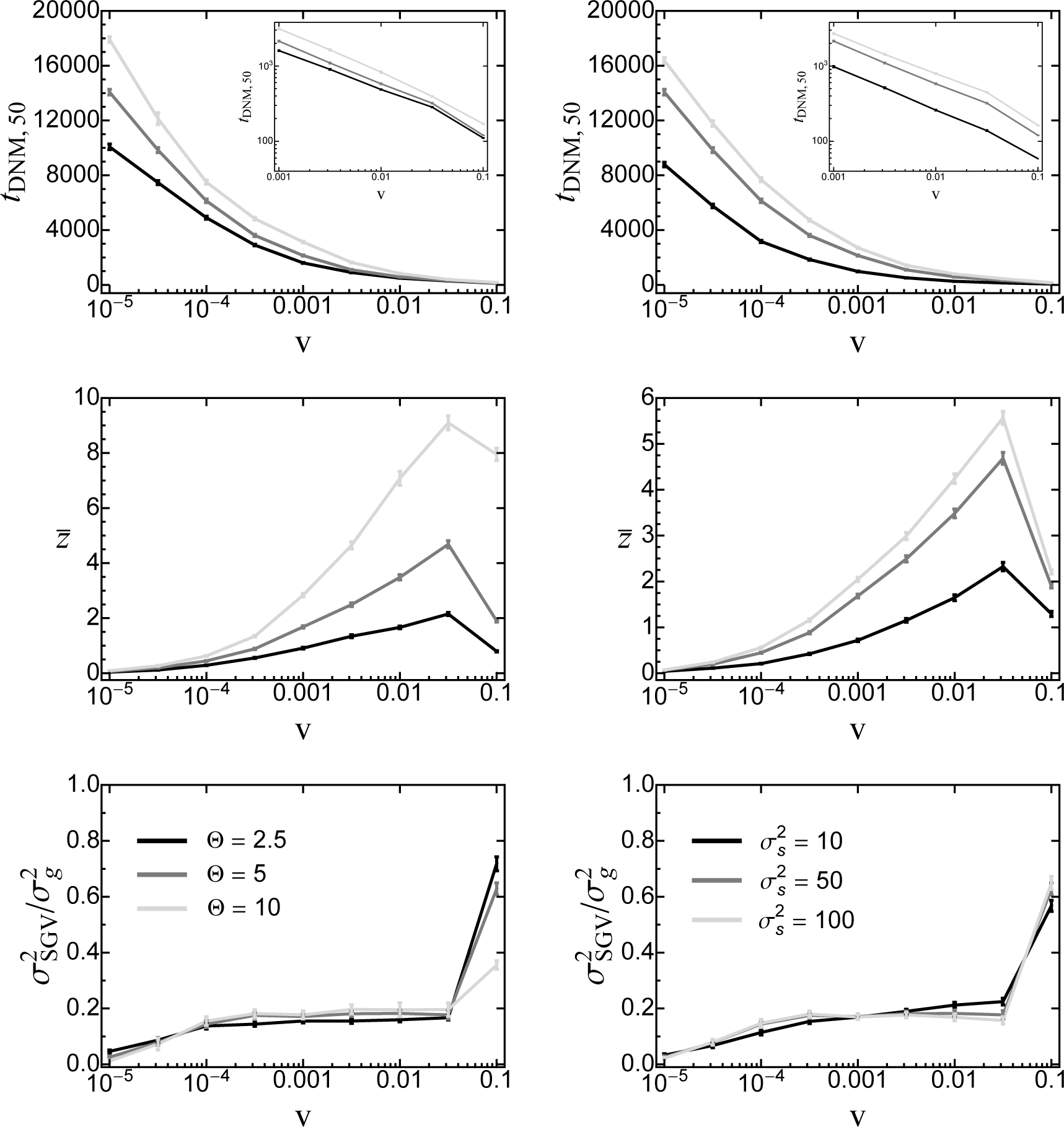
First row: the point in time *t*_DNM,50_ (*z̄*) where 50% of the phenotypic response to moving-optimum selection have been contributed by *de-novo* mutations as a function of the rate of environmental change for various values of Θ (left) and 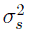 (right). Insets show the results for large *v* on a log-scale. Second row: The mean total phenotypic response at this time. Third row: The relative contribution of original standing genetic variation to the total genetic variance at time *t*_DNM 50_ (*z̄).* Data are means (and standard deviations) from 1000 replicate simulation runs. Fixed parameters (if not stated otherwise): 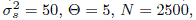 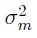 = 0.05.

## DISCUSSION

Global climate change has forced many populations to either go extinct or adapt to the altered environmental condition. When studying the genetic basis of this process, most theoretical work has focused on adaptation from new mutations (e.g., Gillespie 1984; Orr 1998, 2000; Collins *et al.* 2007; Kopp and Hermisson 2007, 2009a,b; Matuszewski *et al.* 2014). Consequently, very little is known about the details of adaptation from standing genetic variation (but see Orr and Betancourt 2001; Hermisson and Pennings 2005), that is, which of the alleles segregating in a population will become fixed and contribute to the evolutionary response. Here, we have used analytical approximations and stochastic simulations to study the effects of standing genetic variation on the genetic basis of adaptation in gradually changing environments. Supporting a verbal hypothesis by Barrett and Schluter (2008), we show that, when comparing adaptation from standing genetic variation to that from *de-novo* mutations, the former proceeds, on average, by the fixation of more alleles of small effect. In both cases, however, the genetic basis of adaptation crucially depends on the efficacy of selection, which in turn is determined by the population size, the strength of (stabilizing) selection and the rate of environmental change. When standing genetic variation is the sole source for adaptation, we find that fast environmental change enables the population to traverse larger distances in phenotype space than slow environmental change, in contrast to studies that consider adaptation from new mutations only (Perron *et al.* 2008; Bell and Gonzalez 2011; Lindsey *et al.* 2013; Bell 2013). We now discuss these results in greater detail.

### The genetic basis of adaptation in the moving-optimum model

Introduced as a model for sustained environmental change, such as global warming (Lynch *et al.* 1991; Lynch and Lande 1993), the moving-optimum model describes the evolution of a quantitative trait under stabilizing selection towards a time-dependent optimum (Bürger 2000). A large number of studies have analyzed both the basic model and several modifi-cations, for example, models with a periodic or fluctuating optimum, or models for multiple traits (Slatkin and Lande 1976; Charlesworth 1993; Bürger and Lynch 1995; Lande and Shannon 1996; Kopp and Hermisson 2007, 2009a,b; Gomulkiewicz and Houle 2009; Zhang 2012; Chevin 2013; Matuszewski *et al.* 2014). Following traditional quantitative-genetic approaches, the majority of these studies assumed that the distribution of genotypes (and phenotypes) is Gaussian with constant (time-invariant) genetic variance, and they have mostly focussed on the evolution of the population mean phenotype and on the conditions for population persistence (Bürger and Lynch 1995; Lande and Shannon 1996; Gomulkiewicz and Houle 2009). None of these models, however, allows to address the fate of individual alleles (i.e., whether they become fixed or not). In a recent series of papers on the moving-optimum model, Kopp and Hermisson (2007, 2009a,b) studied the genetic basis of adaptation from new mutations and derived the distribution of adaptive substitutions (i.e, the distribution of the phenotypic effects of those mutations that arise and become fixed in a population); this approach has recently been generalized to multiple phenotypic traits by Matuszewski *et al.* (2014). The shape of this distribution resembles a Gamma-distribution with an intermediate mode. Thus, most substitutions are of intermediate effect with only a few large-effect alleles contributing to adaptation. The reason is that small-effect alleles - despite appearing more frequently than large-effect alleles - have only small effects on fitness (and are, hence, often lost due to genetic drift), while large-effect alleles might be removed because they “overshoot” the optimum (Kopp and Hermisson (2009b)). A detailed comparison and discussion of the distribution of adaptive substitutions from *de-novo* mutations with (eq. 23) and without (Kopp and Hermisson (2009b)) genetic background variation is given in Supporting Information 2.

Here, we have studied the genetic basis of adaptation from standing genetic variation. We find that the distribution of substitutions from standing genetic variation depends on the distribution of standing genetic variants (i.e., distribution of alleles segregating in the population prior to the environmental change) and the intensity of selection. The former is shaped primarily by the distribution of new mutations and the strength of stabilizing selection, which removes large-effect alleles. Depending on the speed of change *v*, we find two regimes that are characterized by separate distributions of standing substitutions. If the environment changes sufficiently fast, the distribution of adaptive substitutions resembles a lognormal distribution with a strong contribution of small-effect alleles (eq. 19; Fig. 2). The reason is that, in the standing genetic variation, small-effect alleles are more frequent than large-effect alleles and might already segregate at appreciable frequency (so that they are not lost by genetic drift). With a moving optimum, they furthermore are the first to become positively selected, hence reducing the time they are under purifying selection. Finally, epistatic interactions between co-segregating alleles (or between a focal allele and the genetic background) also favor alleles of small effect. Consequently, when adapting from standing genetic variation, most substitutions are of small phenotypic effect.

The second regime occurs if the rate of environmental change *v* is very small. In this case, allele-frequency dynamics are dominated by genetic drift, and the distribution of adaptive substitutions reflects the approximately Gaussian distribution of standing genetic variants (eq. 25; Fig. S3_2). It should be noted, however, that fixations under this regime take a very long time, similar to that of purely neutral substitutions (i.e., on the timescale of 4*N_e_*).

Finally, we have studied the relative importance of standing genetic variation and *de-novo* mutations over the course of adaptation. As shown in Figures 7 and 8, the initial response to selection is almost entirely based on standing variation, with *de-novo* mutations becoming gradually more important. The time scale of this transition strongly depends on the rate of environmental change, but for slow or moderately fast change, it typically occurs over at least hundreds of generations (Figs. 8, S3_8 and Figs. S3_9, S3_10). This observation is in contrast to results by Hill and Rasbash (1986b), who found that under strong artificial (i.e., truncation) selection in small populations (*N* = 20), new mutations might contribute up to one third of the total response after as little as 20 generations. Our results show that the situation is very different for large populations under natural selection in gradually changing environments. The likely reason for this difference is that truncation selection induces strong directional selection (corresponding to large *v*) and only extreme phenotypes reproduce. Thus, truncation selection is much more efficient in maintaining large-effect *de-novo* mutations, while eroding genetic variation more quickly (because it introduces a large skew in the offspring distribution). However, the similarities and differences in the genetic basis of responses to artificial versus natural selection is an interesting topic—in particular, for the interpretation of the large amount of genetic data available from breeding programs (Stern and Orgonzo 2009)—that should be addressed in future studies.

Throughout this study, we have focused on adaptation to a moving optimum, that is, a scenario of gradual environmental change. An obvious question is how our results would change under the alternative scenario of a one-time sudden shift in the optimum (as assumed in numerous studies, e.g., Orr 1998; Hermisson and Pennings 2005; Chevin and Hospital 2008). While beyond the scope of this paper, our approach should, in principle, still be applicable. In particular, each focal allele still experiences a gradual change in its selection coefficient, due to the evolution of the genetic background. Unlike in the moving-optimum model, however, the selection coefficient *decreases*, as the mean phenotype gradually approaches the new optimum. Hence, a suitably modified version of equation 13 would give the probability that a focal allele *establishes* in the population (i.e., escapes stochastic loss), but in the absence of continued environmental change, establishment does not guarantee fixation. In other words, alleles need to “race for fixation” before other competing alleles get fixed and they become deleterious (Kopp and Hermisson 2007, 2009a). The dynamics of a mutation along its trajectory should therefore be even more complex than in the moving-optimum model, and show an even stronger dependence on the genetic background (Chevin and Hospital 2008).

### Extinction and the rate of environmental change

Recently, several experimental studies have explored how the rate of environmental change affects the persistence of populations that rely on new mutations for adapting to a gradually changing environment (Perron *et al.* 2008; Bell and Gonzalez 2011; Lindsey *et al.* 2013). In line with theoretical predictions (Bell 2013), all studies found that “evolutionary rescue” is contingent on a small rate of environmental change. In particular, Lindsey *et al.* (2013) evolved replicate populations of *E. coli* under different rates of increase in antibiotic concentration and found that certain genotypes were evolutionarily inaccessible under rapid environmental change, suggesting that “rapidly deteriorating environments not only limit mutational opportunities by lowering population size, but […] also eliminate sets of mutations as evolutionary options”. This is in stark contrast to our prediction that faster environmental change can enable the population to remain better adapted and to traverse larger distances in phenotype space when standing genetic variation is the sole source for adaptation (Fig. 6 and Figs S3_6, S3_7; in line with recent experimental observations; H. Teotonio, private communication). The difference between these results arises from the availability of the “adaptive material”. While *de-novo* mutations first need to appear and survive stochastic loss before becoming fixed, standing genetic variants are available right away and might already be segregating at appreciable frequency. Thus, in both cases, the rate of environmental change plays a critical, though antagonistic, role in determining the evolutionary options. While fast environmental change eliminates sets of new mutations, it simultaneously helps to preserve standing genetic variation until it can be picked-up by selection. Under slow change, in contrast, most large-effect alleles from the standing variation, by the time they are needed, are already eliminated by drift or stabilizing selection.

Our results also mean that, if the optimum stops moving at a given value *z*_opt,max_, populations will achieve a higher degree of adaptation (higher *z̄*)* if the final optimum is reached fast rather than slowly (see also Uecker and Hermisson 2014), at least if standing genetic variation is the sole source for adaptation. While this assumption is an obvious simplification, it may often be approximately true in natural populations. The same holds true in experimental populations, where selection is usually strong and the duration of the experiment short, such that *de-novo* mutations can frequently be neglected (see Fig. 8).

### Testing the predictions

The predictions made by our model can in principle be tested empirically, even though suitable data might be sparse and experiments challenging. There is, of course, ample evidence for adaptation from standing genetic variation. For example, Domingues *et al.* (2012) showed that camouflaging pigmentation of oldfield mice (*Peromyscus polionotus)* that have colonized Florida’s Gulf Coast has evolved quite rapidly from a pre-existing mutation in the *Mclr* gene; Limborg *et al.* (2014) investigated selection in two allochronic but sympatric lineages of pink salmon (*Oncorhynchus gorbuscha)* and identified 24 divergent loci that had arisen from different pools of standing genetic variation, and Turchin *et al.* (2012) showed that height-associated alleles in humans display a clear signal for widespread selection on standing genetic variation.

However, testing the predictions of our model requires, in addition, detailed knowledge of the genotype-phenotype relation. Currently, there is only a small (yet increasing) number of systems for which both a set of functionally validated beneficial mutations and their selection coefficients under different environmental conditions are available (Jensen 2014). Thus, estimating the distribution of standing substitutions will be challenging, because of the often unknown phenotypic and fitness effects of beneficial mutations and the large number of replicate experiments needed to obtain a reliable empirical distribution. Furthermore, even if these problems were solved, small-effect alleles might not be detectable due to statistical limitations (Otto and Jones 2000), and in certain limiting cases where the population quickly goes extinct (i.e., when the environment changes very fast), the distribution of adaptive substitutions from standing genetic variation might be indistinguishable to that from *de-novo* substitutions (Fig. 5).

Recent developments in laboratory systems (Morran *et al.* 2009; Parts *et al.* 2011), however, have created opportunities for experimental evolution studies in which population size,the selective regime and the duration of selection can be manipulated, and adaptation from *de-novo* mutations and standing genetic variation can be recorded (Burke 2012). Applying these techniques in experiments in the vein of Lindsey *et al.* (2013), but starting from a polymorphic population, should make it possible to test the relation between the rate of environmental change and population persistence, and to assess the probability of adaptation from standing genetic variation. First experiments along these lines are currently being carried out in populations of *C. elegans*, with the aim of determining the limits of adaptation to different rates of increase in sodium chloride concentration (H. Teotonio, private communication). Furthermore, Pennings (2012) recently applied the Hermisson and Pennings (2005) framework to show that standing genetic variation plays an important role in the evolution of drug-resistance in *HIV*, affecting up to 39% of patients (depending on treatment) and explaining why resistance mutations in patients who interrupt treatment are likely to become established within the first year. A similar approach should also be applicable to scenarios of gradual environmental change (e.g., evolution of resistance mutations under gradually increasing antibiotic concentrations).

## Conclusion

As global climate change continues to force populations to respond to the altered environmental conditions, studying adaptation to changing environments - both empirically and theoretically - has become one of the main topics in evolutionary biology. Despite increased efforts, however, very little is known about the genetic basis of adaptation from standing genetic variation. Our analysis of the moving-optimum model shows that this process has, indeed, a very different genetic basis than that of adaptation from *de-novo* mutations. In particular, adaptation proceeds via the fixation many small-effect alleles (and just a few large ones). In accordance with previous studies, the adaptive process critically depends on the tempo of environmental change. Specifically, when populations adapt from standing genetic variation only, the potential for adaptation increases as the environment changes faster.

## Acknowledgements

We thank R. Bürger, LM. Chevin, and C. Vogl for constructive comments on this manuscript. This study was supported by Austrian Science Fund, FWF (grant P 22581-B17 to MK and grant P22188 to Reinhard Bürger), Austrian Agency for International Cooperation in Education and Research, OEAD (grant FR06/2014 to JH), Campus France (grant PHC AMADEUS 31642SJ to MK), and a Writing-Up Fellowship from the Konrad Lorenz Institute for Evolution and Cognition Research (KLI) to SM.

### APPENDIX

#### Appendix 1: Theoretical Background

In this Appendix, we briefly recapitulate results from previous studies that form the basis for our analytical derivations.

#### The probability of adaptation from standing genetic variation for a single biallelic locus after a sudden environmental change

Hermisson and Pennings (2005) studied the situation where the selection scheme at a single bi-allelic locus changes following a sudden environmental change. In particular, they derived the probability for a mutant allele to reach fixation that was neutral or deleterious prior to the change but has become beneficial in the new environment. In the continuum limit for allele frequencies this probability is given by

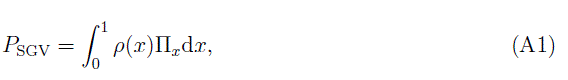

where *ρ(x)* is the density function for the allele frequency *x* of the mutant allele in mutation-selection-drift balance and Π_*χ*_ denotes its fixation probability.

For a mutant allele present at frequency *x* and with selective advantage *s*_*b*_ in the new environment, the fixation probability is given by (Kimura 1957)

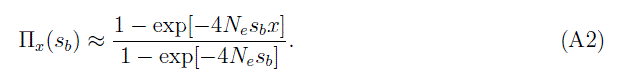

There are two points to make here. First, mutational effects in the Hermisson and Pennings (2005) model are directly proportional to fitness, whereas mutations in our model affect a phenotype under selection. Second, in our framework, *s*_*b*_ denotes the (beneficial)selection coefficient for heterozygotes.

Approximations for *ρ(x)* can be derived from standard diffusion theory (Ewens 2004; for details see Hermisson and Pennings 2005). If the mutant allele was neutral prior to the change in the selection scheme

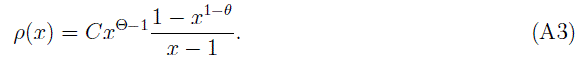

Here, *C* = (*γ* + *ψ*(θ))^−1^ denotes a normalization constant where γ ≈ 0.577 is Euler’s gamma and *ψ*(·) is the polygamma function. Similarly, if the mutant allele was deleterious before the environmental change (with negative selection coefficient *s_d_*) the allele-frequency distribution is given by

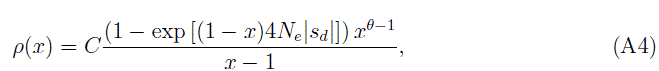

where *C* = (_1_*F*_1_(0,θ, 4 *N*_*e*_|*s_d_*|))^−1^ denotes a normalization constant and *_1_F_1_*(*a,b,c)* is the hypergeometric function. If the allele was sufficiently deleterious (4 *N_e_*|*s_d_*| ≥ 10), equation (A4) can further be approximated as

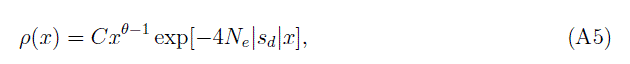

where 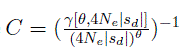 again denotes a normalization constant with 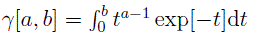 denoting the lower incomplete gamma function.

Finally, the probability that a population successfully adapts from standing genetic variation can be derived as

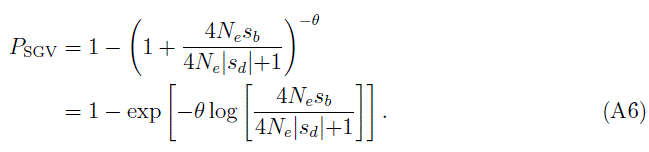

#### Fixation probabilities under time-inhomogeneous selection

In gradually changing environments, the selection coefficient of a given (mutant) allele is not fixed but changes over time (i.e., as the position of the optimum changes). Uecker and Hermisson (2011) recently developed a mathematical framework based on branching-process theory to describe the fixation process of a beneficial allele under temporal variation in population size and selection pressures. They showed that the probability of fixation of a mutation starting with *n* initial copies is given by

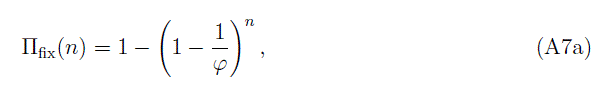

where

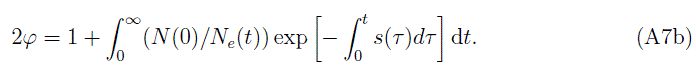

Assuming that the population size remains constant and that the selection coefficient increases linearly in
time, *s(t) = s*_*d*_ +*s_v_t*, equation (A7a) becomes

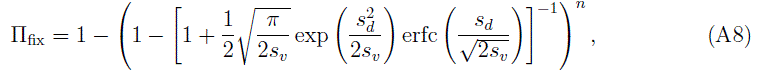

where erfc(·) denotes the complementary Gaussian error function.

### SUPPORTING INFORMATION

#### Supporting Information 1: Limited Recombination

**Figure S1_1.**
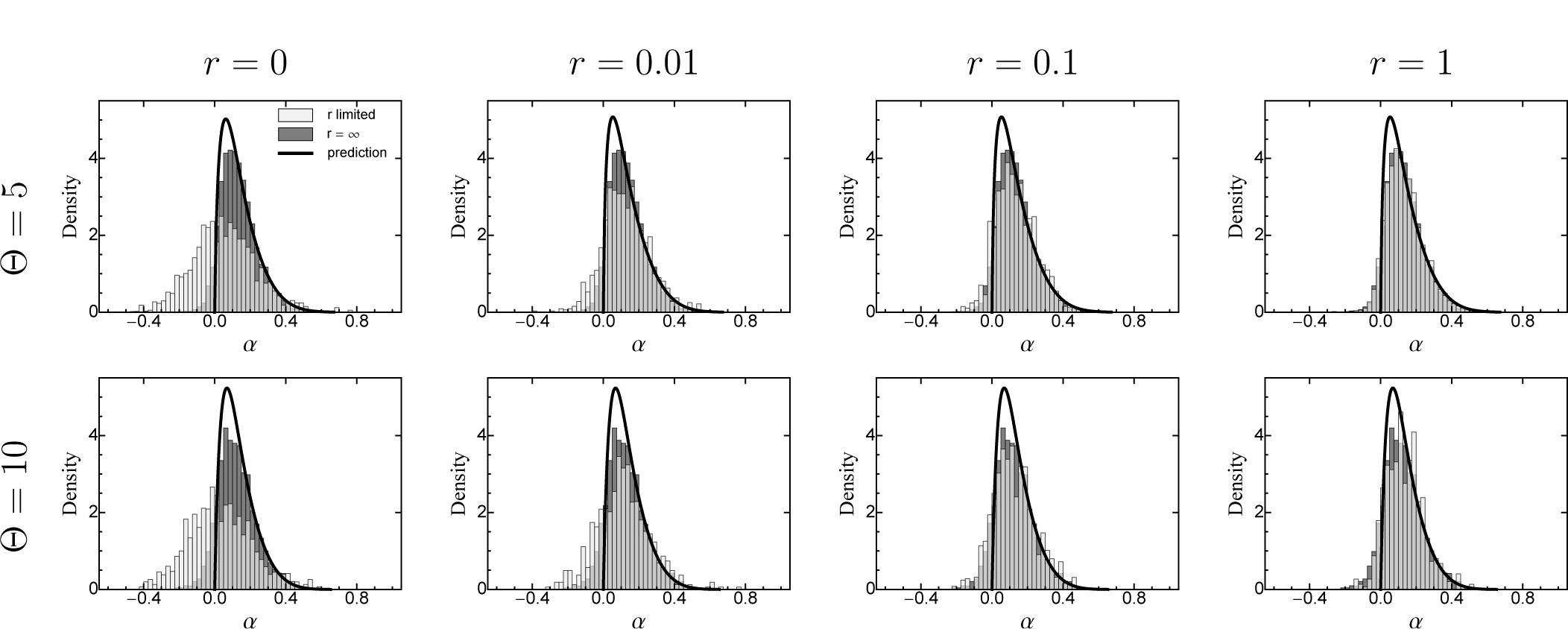
The distribution of adaptive substitutions from standing genetic variation for free recombination (dark bins) compared to that for limited recombination (light bins). The black line corresponds to the analytical prediction (eq. 19). 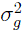 is given by equation (16). Fixed parameters: 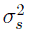 = 50, *N* = 2500, *v* = 0.001, 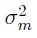 = 0.05.

**Figure S1_2.**
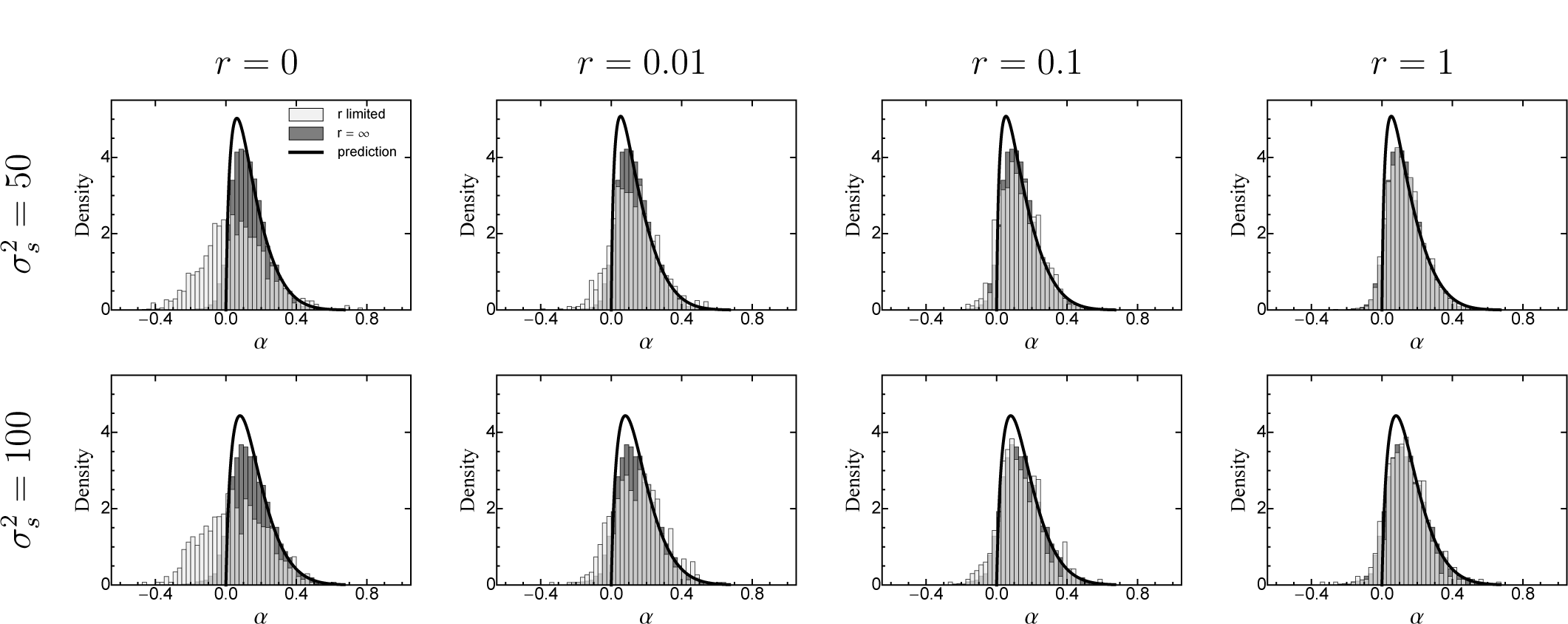
The distribution of adaptive substitutions from standing genetic variation for free recombination (dark bins) compared to that for limited recombination (light bins). The black line corresponds to the analytical prediction (eq. 19). 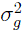 is given by equation (16). Fixed parameters: Θ = 5, *N* = 2500, *v* = 0.001,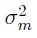 = 0.005.

**Figure S1_3.**
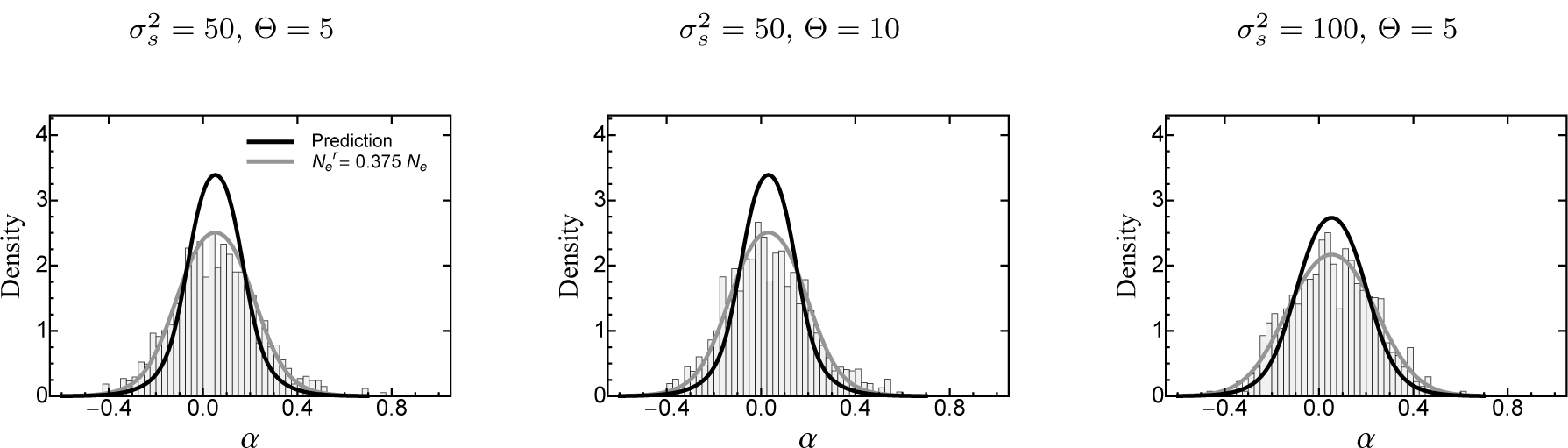
The distribution of adaptive substitutions from standing genetic variation for complete linkage (no recombination). The black and the grey line corresponds to the analytical prediction (eq. 25) that are centred around the mean of the individual-based simulation. For the grey line Ne has been adjusted by a factor 0.385 to match the distribution from the individual-based simulations. Other parameters: *v* = 0.001, *r* = 0, *N* = 2500, 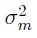 = 0.005.

The individual-based simulation results presented in the main text were obtained under the assumption of free recombination. In this Supplementary Information, we relax this assumption and study the effects of linkage (i.e., limited recombination).

We first clarify the meaning of the recombination parameter *r*, which determines the mean number of crossover events per meiosis. By definition, the simulated genome corresponds to a single chromosome of length *D_𝒢_ = r ·* 100cM, and the mean distance between two randomly chosen sites is 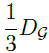. The mean distance between two adjacent polymorphic loci is 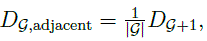, where 𝒢 is the mean number of polymorphic loci, which depends on Θ and 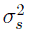 (eq. 26).

The corresponding recombination rate **τ** between two polymorphic loci is given by the inverse of Haldane’s mapping function (Speed 2005), that is,

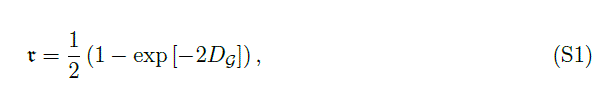

see Table S1_1.

**Table S1_1.**
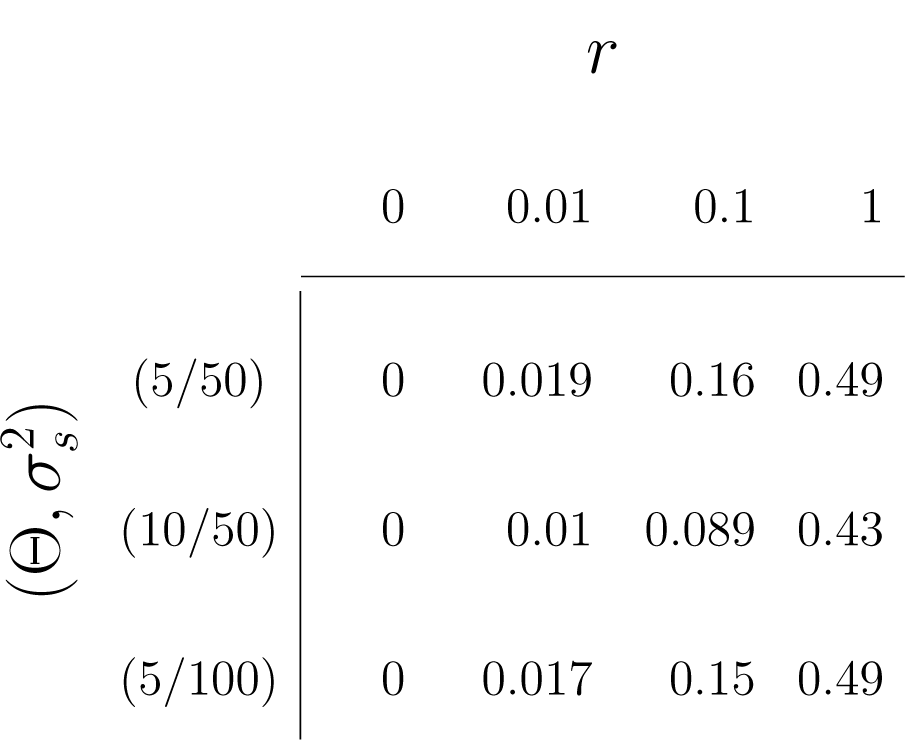
The classical population genetic recombination rate **τ** (eq. S1) between two adjacent loci for different values of 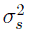, Θ and *r*. Other parameters: 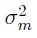 = 0.05.

The effect of limited recombination on the distribution of adaptive substitutions from standing genetic variation is illustrated in Figures S1_1 and S1_2. For *r* = 1 (corresponding to a genome length of 100cM and an average recombination rate *r* of close to 0.5, see table S1_1), the distribution is essentially identical to that for linkage equilibrium. As *r* decreases, the distribution progressively shifts to the left, becomes more symmetric and includes more and more alleles with negative phenotypic effect. For *r* = 0 (corresponding to complete linkage or asexual reproduction), it resembles the distribution for “drift-driven” evolution (i.e., when selection is not efficient; Fig. S3_2). The reason is that fixation involves entire haplotypes carrying multiple mutations, whose (positive and negative) effects largely cancel. From a different perspective, limited recombination leads to Hill-Robertson interference between cosegregating alleles (Hill and Robertson 1966), which in many respects corresponds to a decrease in effective population size *N*_*e*_ (Comeron *et al.* 2008), which in turn reduces the efficacy of selection. Note, however, that unlike in the case of a slowly changing environment (Fig. S3_2) reducing *N*_*e*_ also affects the equilibrium allele-frequency distribution *ρ(x, α)* (by reducing the strength of selection against large-effect alleles). In line with previous simulation results (Comeron *et al.* 2008), we find that equation (25) provides a very good fit, when *N*_*e*_ is set to 38.5% of its original value.

#### Supporting Information 2 The distribution of adaptive substitutions from *de-novo* mutations with and without genetic background variation

There are two ways in which the distribution of adaptive substitutions from standing genetic variation can be compared to that from *de-novo* mutations. The first comparison considers a population without genetic background variation. This is the situation studied by Kopp and Hermisson (2009a), where an essentially monomorphic population performs an adaptive walk following a moving optimum. The second situation is the one described by equation (23), where new mutations interact with a genetic background of constant variance (this background is presumably itself constantly replenished by new mutations). Analytical predictions for all three distributions are compared in Figures. S2_1, S2_2 and S2_3. It can be seen that the adaptive-walk prediction (eq. 14 in Kopp and Hermisson (2009b); red line) is always shifted towards larger α compared to the distribution of adaptive substitutions from standing genetic variation (eq. 19, black line). The predicted distribution from *de-novo* mutations in the presence of genetic background variation (eq. 23, grey curve) shifts from the latter to the former as *v* increases. The reason is that, for small *v*, the fixation of both standing variants and new mutations in the presence of background variation is strongly constrained by the equilibrium lag (eq. 9). For large v, in contrast, the lag is large and adaptation is primarily limited by the available alleles, independent of their source and initial frequency (mutation-limited regime *sensu* Kopp and Hermisson (2009b)). Note, however, that in both limiting cases, equation (19) is a poor predictor for the simulated substitutions from standing variation (Fig. 5, S3_2). Nevertheless, it remains true that adaptive substitutions from new mutations are generally smaller than those from new mutations, with or without genetic background variation.

**Figure S2_1.**
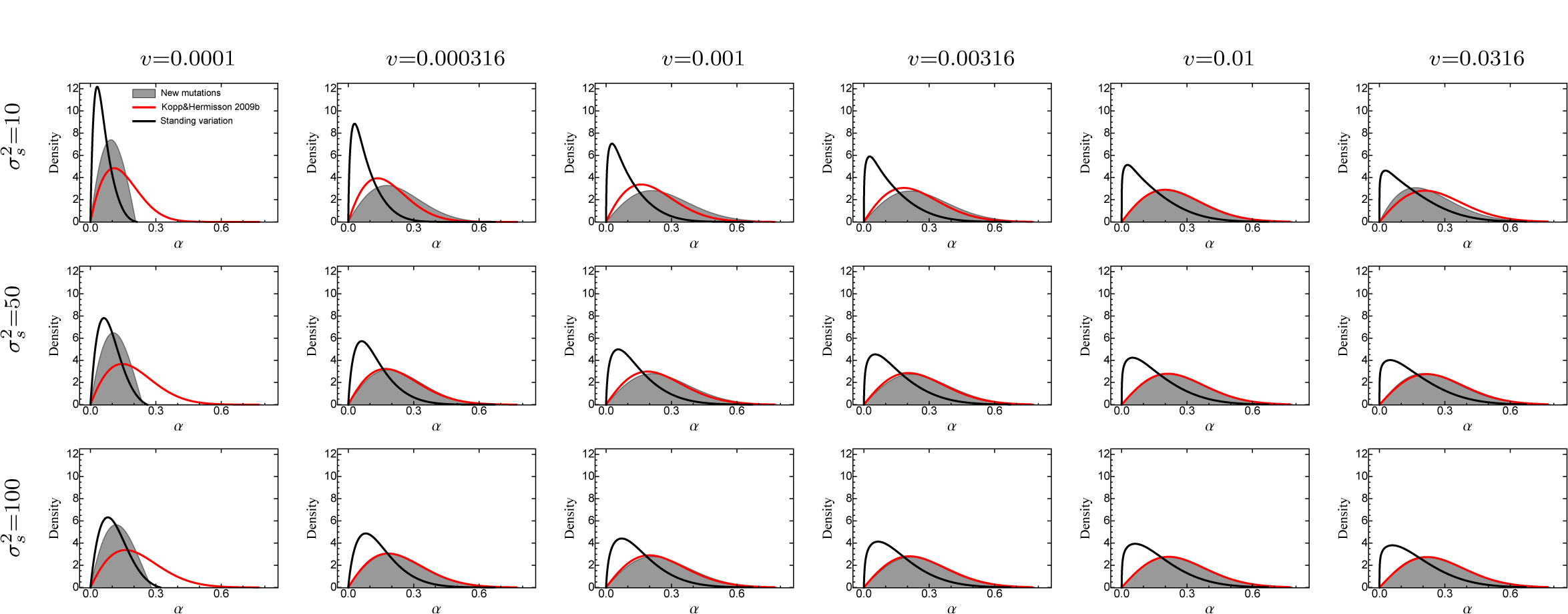
Comparison of the analytical predictions for the distribution of adaptive substitutions from standing genetic variation and de-novo mutations. The black line corresponds to eq. (19), with the genetic background variation 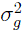 determined by the SHC approximation (eq. 16). The grey curve gives the analytical prediction for substitutions from de-novo mutations under the assumption that the phenotypic lag δ_eq_ has reached an equilibrium (eq. 23). The red line gives the analytical prediction for the first substitution from de-novo mutations under the adaptive-walk assumption that there is no genetic background variation (Kopp and Hermisson (2009b), eq. 14). Note that, for some parameter combinations, the simulated distribution from standing variation deviates from eq. (19). In particular, for small *v*, it approaches the “neutral” prediction eq. (25, see Fig. 3, and for large *v*, it may approach the distribution from new mutations, eq. (23), see Fig. 5. Fixed parameters: Θ = 2.5, *N* = 2500, 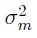 = 0.05.

**Figure S2_2.**
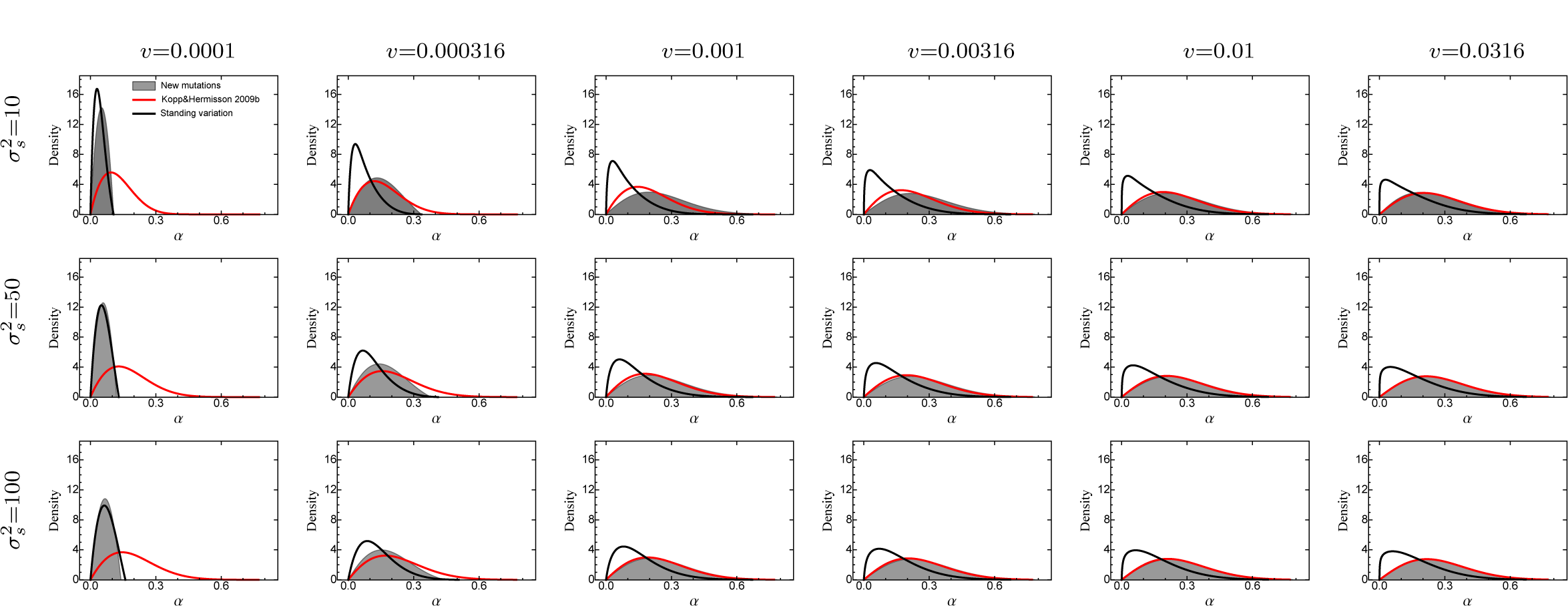
Comparison of the analytical predictions for the distribution of adaptive substitutions from standing genetic variation and de-novo mutations. For further details see Fig. S2_1. Fixed parameters: Θ = 5, *N*2500, 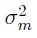 = 0.05.

**Figure S2_3.**
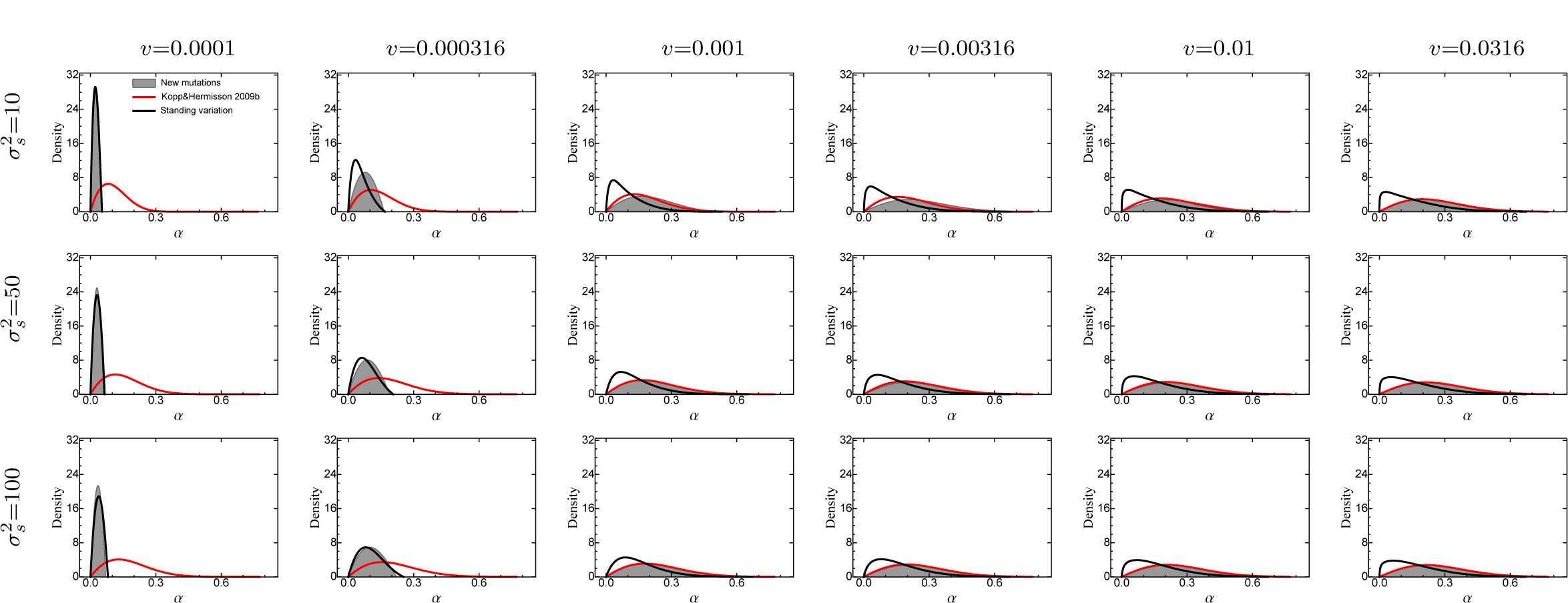
Comparison of the analytical predictions for the distribution of adaptive substitutions from standing genetic variation and de-novo mutations. For further details see Fig. S2_1. Fixed parameters: Θ = 10, *N*2500, 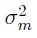 = 0.005.

#### Supporting Information 3: Supporting Figures

**Figure S3_1.**
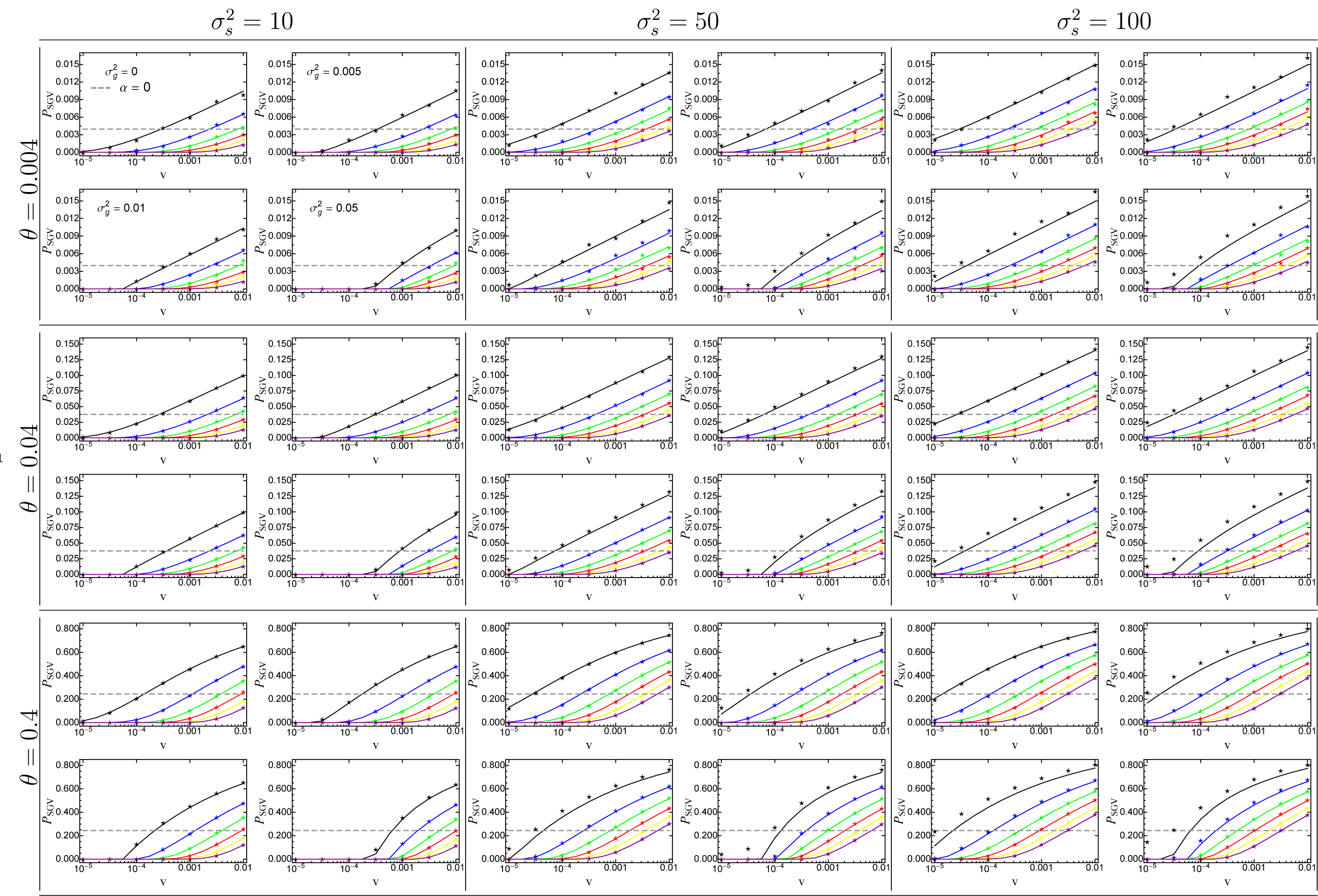
The probability for a mutant allele to adapt from standing genetic variation as a function of the rate of environmental change *v*. Solid lines correspond to the analytical prediction (eq. 14), the grey dashed line shows the probability for a neutral allele (α = 0; eq. 15), and symbols give results from Wright-Fisher simulations. The phenotypic effect size α of the mutant allele ranges from 0.5σ_m_ (top line; black) to 3σ_m_ (bottom line; purple) with increments of 0.5σ_*m*_. The figures in each parameter box (per locus mutation rate θ, width of fitness landscape 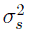) correspond to different values of the genetic background variation 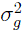 with 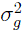 = 0 (no background variation; top left), 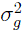 = 0.005. (top right), 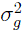 = 0.01 (bottom left) and 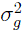 = 0.05 (bottom right). Other parameters: *N_e_* = 25000, 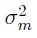 = 0.05.

**Figure S3_2.**
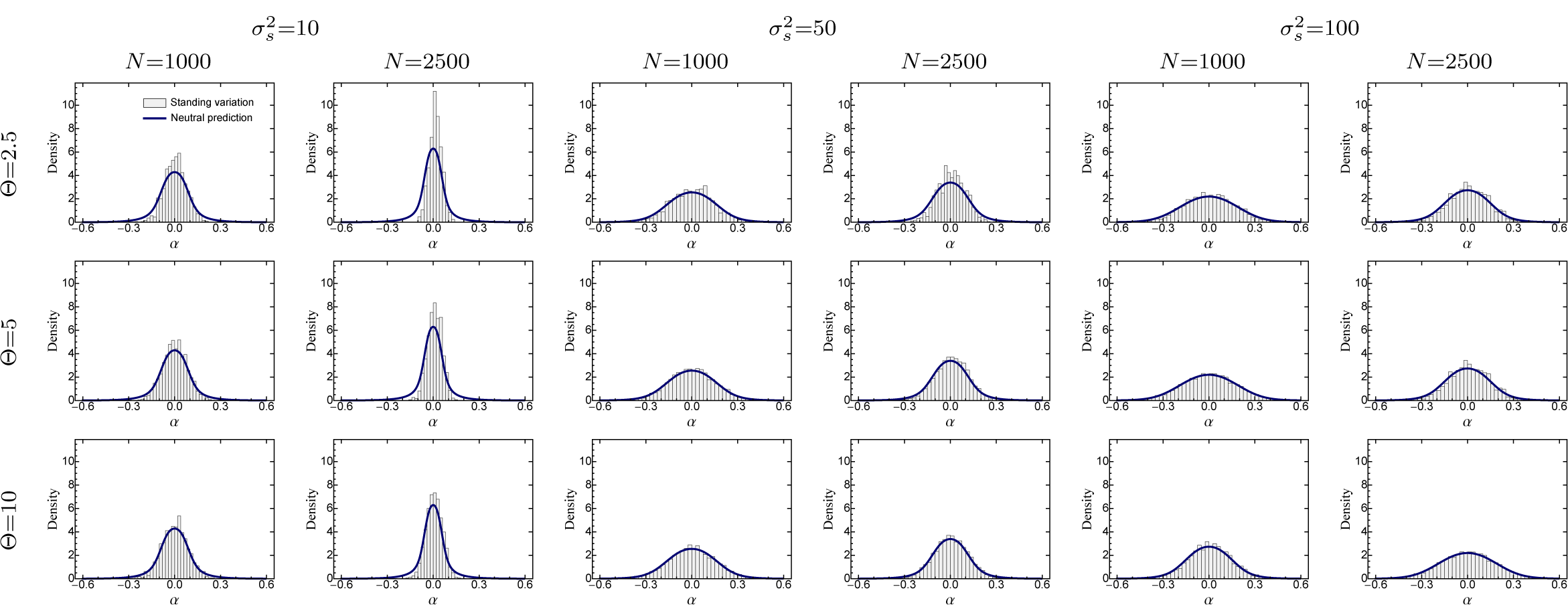
The distribution of adaptive substitutions from standing genetic variation in the case of slow environmental change (*v* = 10^−5^). Histograms show results from individual-based simulations. The blue line gives the analytical prediction (eq. 25), with 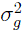 given by eq. 16), which assumes a neutral fixation probability. Fixed parameters: 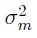 = 0.05.

**Figure S3_3.**
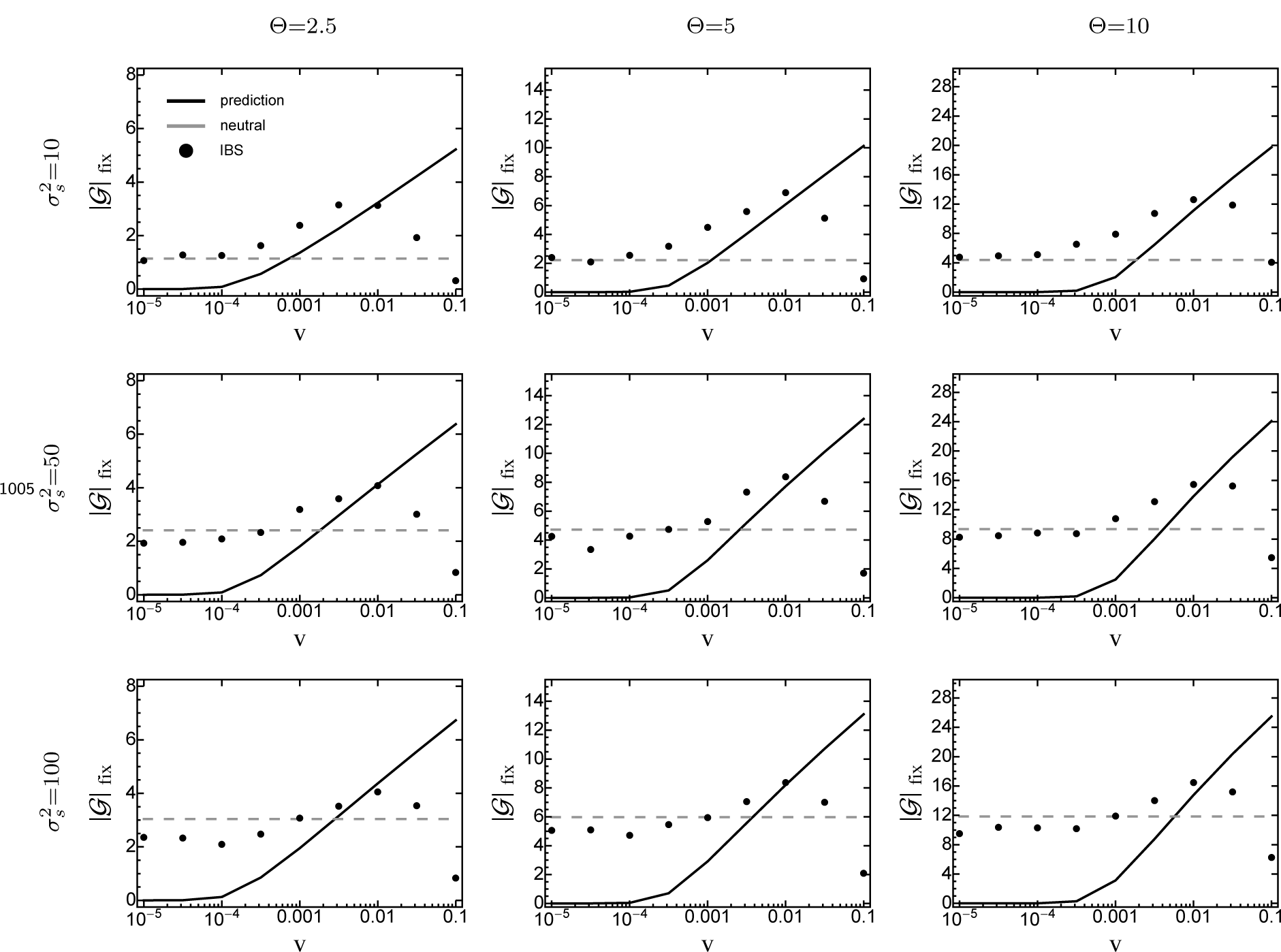
The average number of fixed adaptive substitutions from standing genetic variation, |𝒢|_fix_, as a function of the rate of environmental change *v*, when standing genetic variation is the sole source for adaptation. Symbols show results from individual-based simulations (averaged over 100 replicate runs). The black line gives the analytical prediction (eq. 27) and the grey line corresponds to the average number of neutral fixations 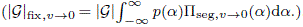. In both cases, 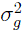 was taken from equation (16). Error bars for standard errors are contained within the symbols. Fixed parameters: *N* = 1000, 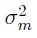 = 0.05.

**Figure S3_4.**
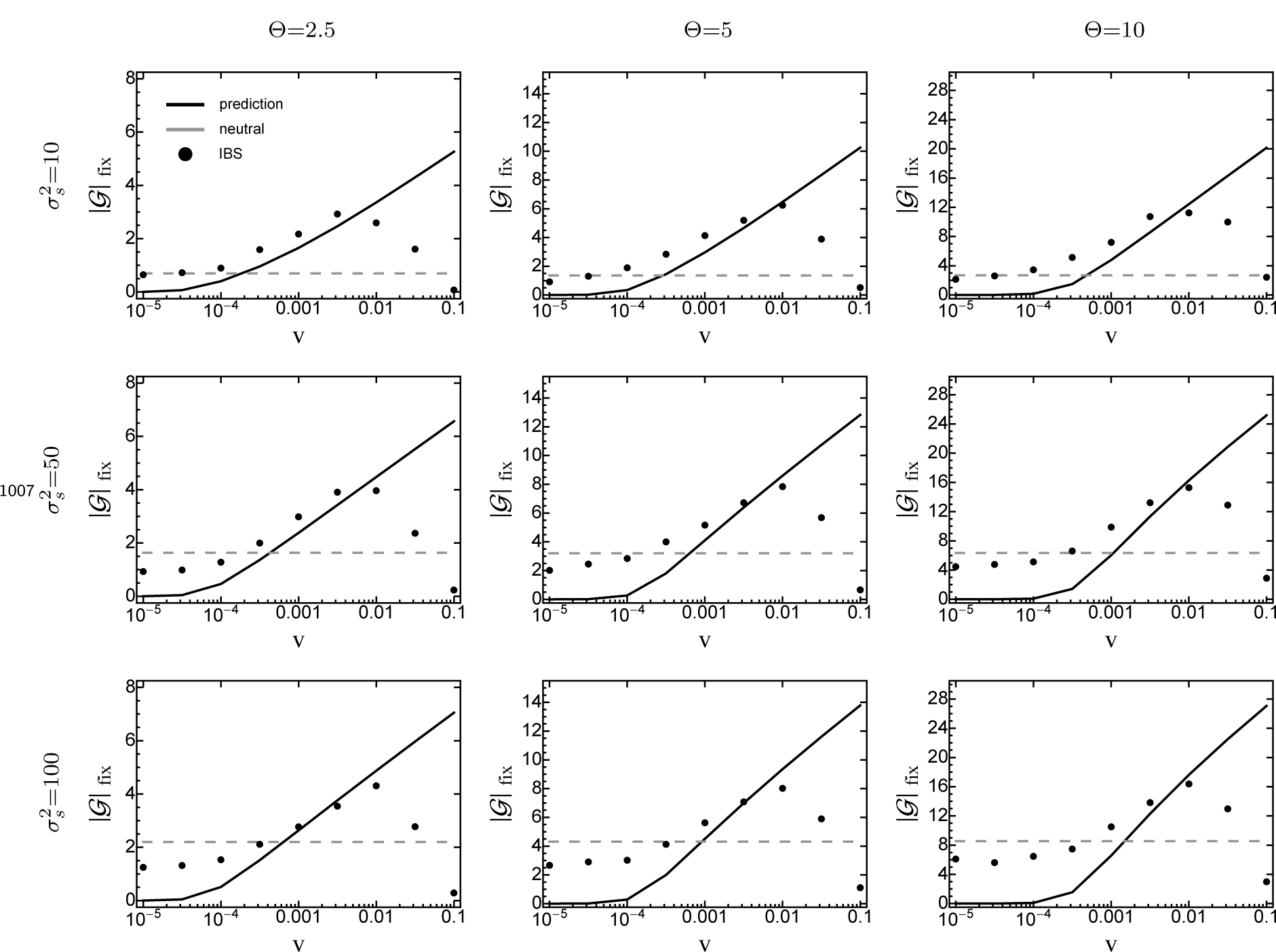
The average number of fixed adaptive substitutions from standing genetic variation, |𝒢|_fix_, as a function of the rate of environmental change *v*, when standing genetic variation is the sole source for adaptation. Symbols show results from individual-based simulations (averaged over 100 replicate runs). The black line gives the analytical prediction (eq. 27) and the grey line corresponds to the average number of neutral fixations 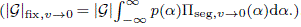. In both cases, 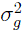 was taken from equation (16). Error bars for standard errors are contained within the symbols. Fixed parameters: *N* = 2500, 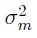 = 0.05.

**Figure S3_5.**
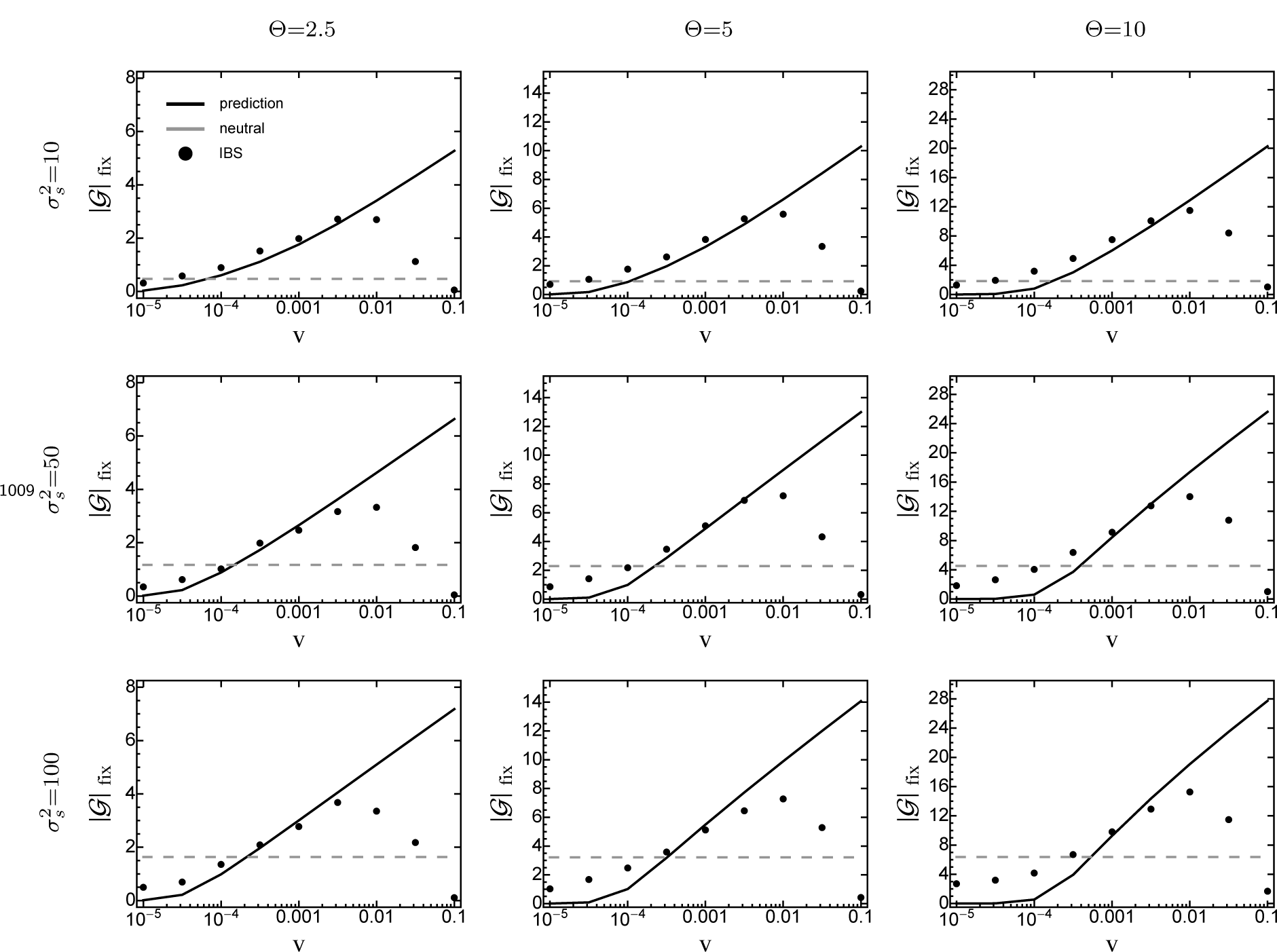
The average number of fixed adaptive substitutions from standing genetic variation, |𝒢|_fix_, as a function of the rate of environmental change *v*, when standing genetic variation is the sole source for adaptation. Symbols show results from individual-based simulations (averaged over 100 replicate runs). The black line gives the analytical prediction (eq. 27) and the grey line corresponds to the average number of neutral fixations 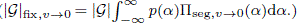. In both cases, 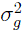 was taken from equation (16). Error bars for standard errors are contained within the symbols. Fixed parameters: *N* = 5000, 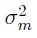 = 0.05.

**Figure S3_6.**
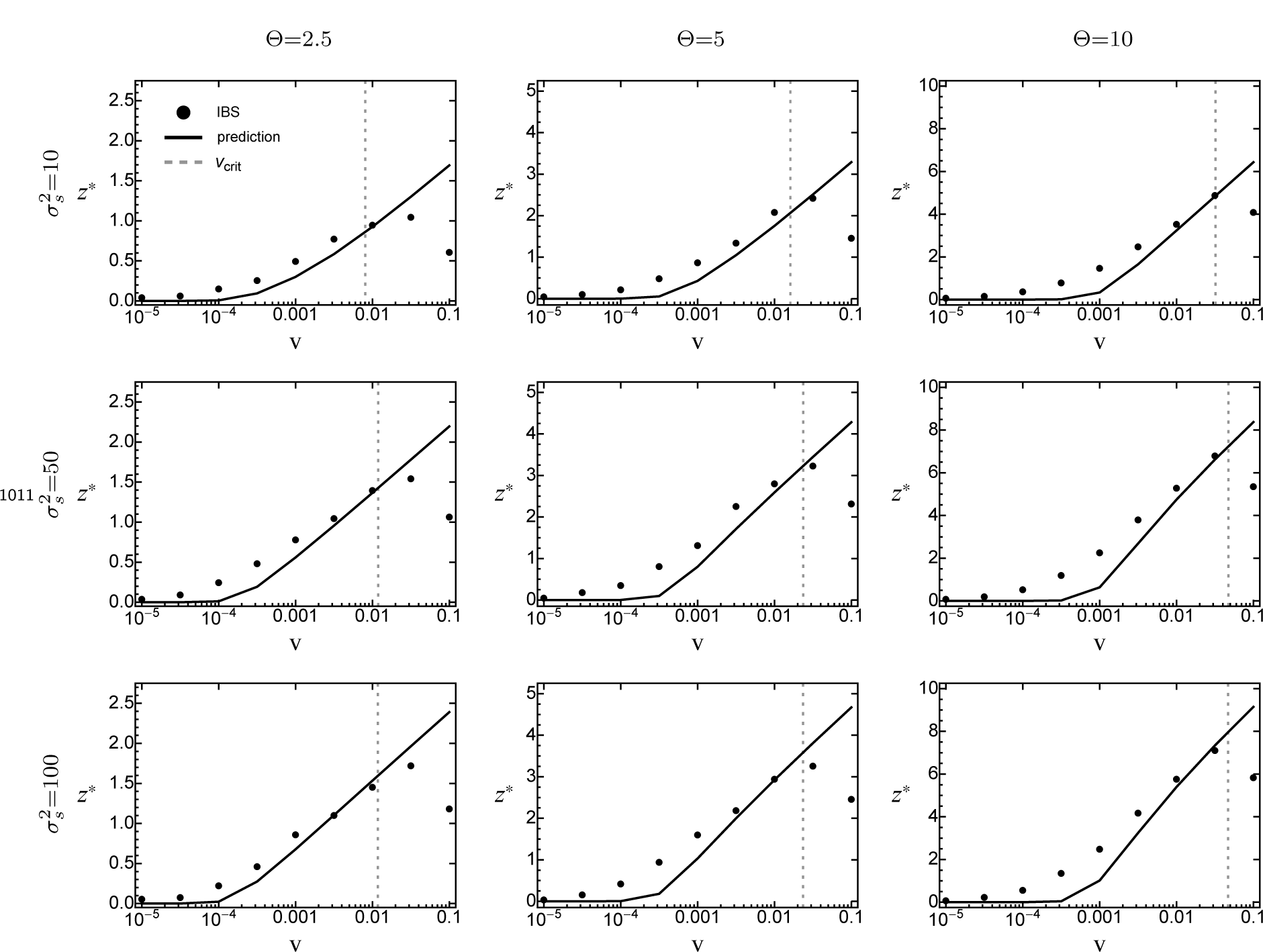
The average distance traversed in phenotype space, *z*^*^, as a function of the rate of environmental change *v*, when standing genetic variation is the sole source for adaptation. Symbols show results from individual-based simulations (averaged over 100 replicate runs). The black line gives the analytical prediction (eq. 28), with 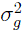 taken from equation (16). The grey-dashed line gives the critical rate of environmental change (eq. 29). Error bars for standard errors are contained within the symbols. Fixed parameters: *N* = 1000, 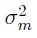 = 0.05.

**Figure S3_7.**
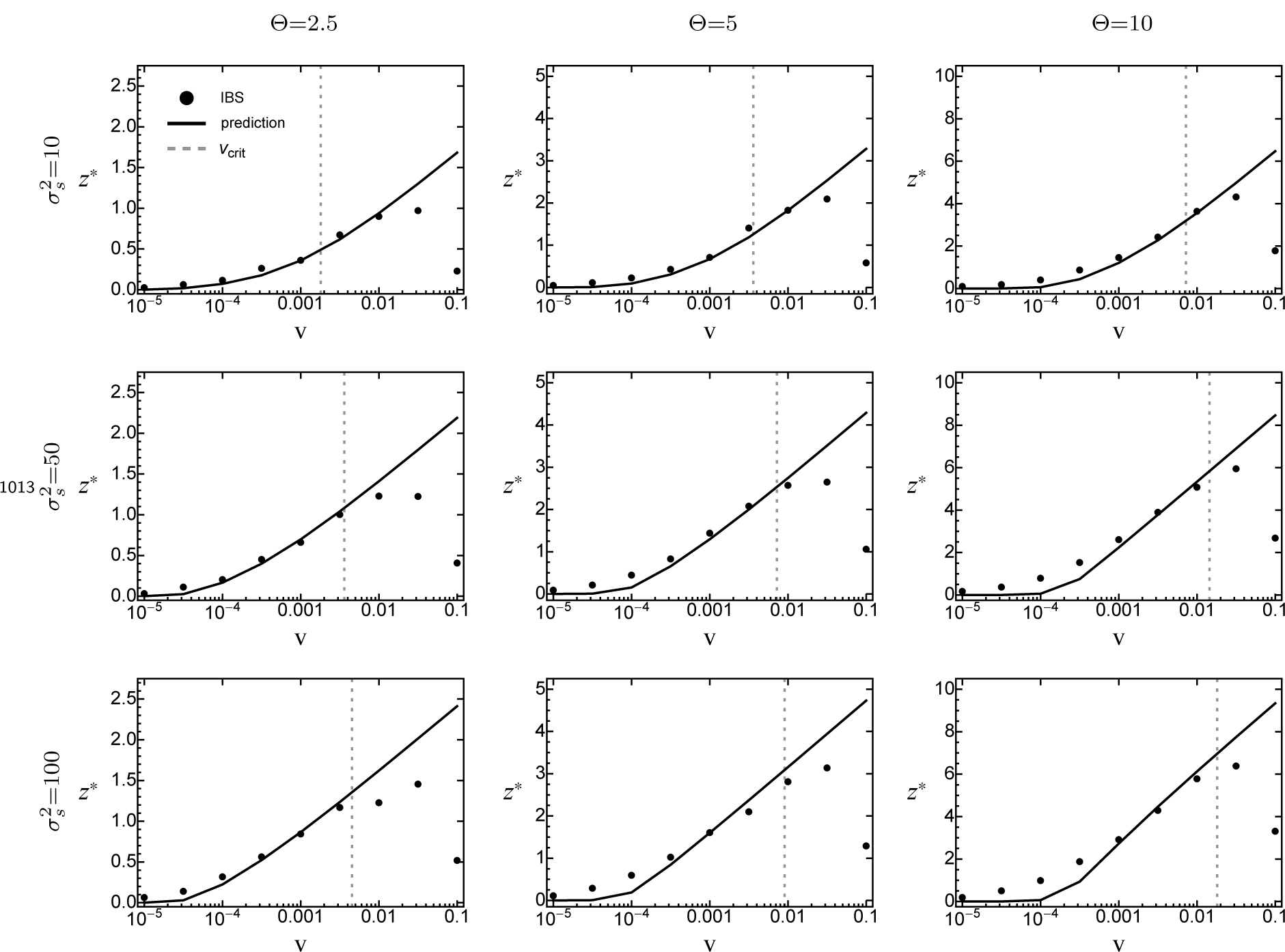
The average distance traversed in phenotype space, *z*^*^, as a function of the rate of environmental change *v*, when standing genetic variation is the sole source for adaptation. Symbols show results from individual-based simulations (averaged over 100 replicate runs). The black line gives the analytical prediction (eq. 28), with 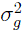 taken from equation (16). The grey-dashed line gives the critical rate of environmental change (eq. 29). Error bars for standard errors are contained within the symbols. Fixed parameters: *N* = 5000, 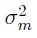 = 0.05.

**Figure S3_8.**
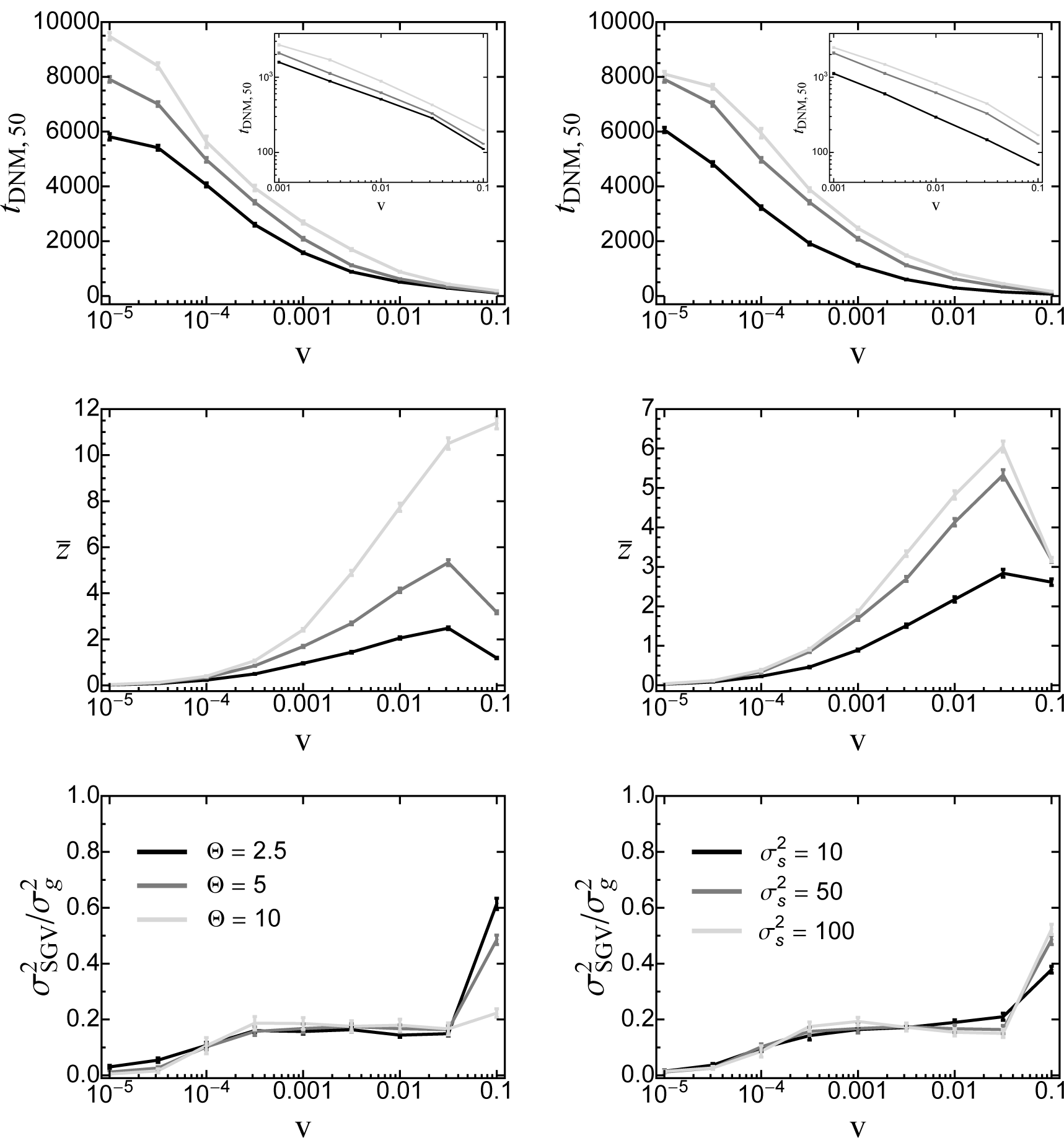
First row: the point in time *t*_DNM,50_ (z̄) where 50% of the phenotypic response to moving-optimum selection have been contributed by de-novo mutations as a function of the rate of environmental change for various values of Θ (left) and 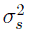 (right). Insets show the results for large *v* on a log-scale. Second row: The mean total phenotypic response at this time. Third row: The relative contribution of original standing genetic variation to the total genetic variance at time *t*_DNM,50_ (z̄). Data are means and standard errors from 1000 replicate simulation runs. Fixed parameters (if not stated otherwise): 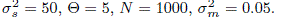

**Figure S3_9.**
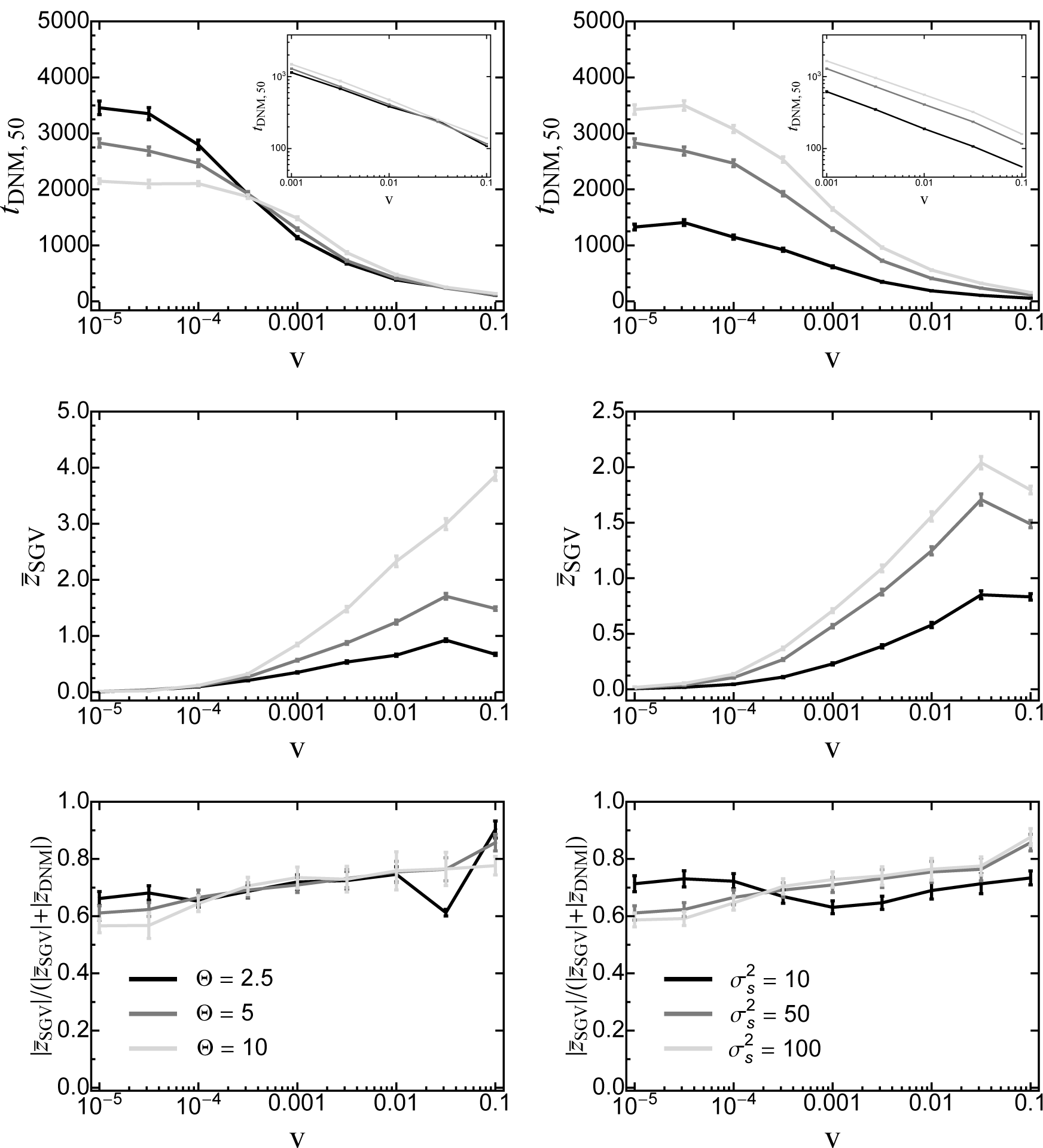
First row: the point in time *t*_DNM,50_ (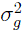) where 50% of the genetic variance is composed of de-novo mutations as a function of the rate of environmental change for various values of Θ (left) and 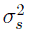 (right). Insets show the results for large *v* on a log-scale. Second row: The mean total phenotypic response from standing genetic variation at this time. Third row: The relative contribution of original standing genetic variation to the total genetic variance at time *t*_DNM,50_ 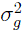. Data are means and standard errors from 1000 replicate simulation runs. Fixed parameters (if not stated otherwise): 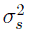 = 50, Θ = 5, *N* = 1000, 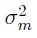 = 0.05.

**Figure S3_10.**
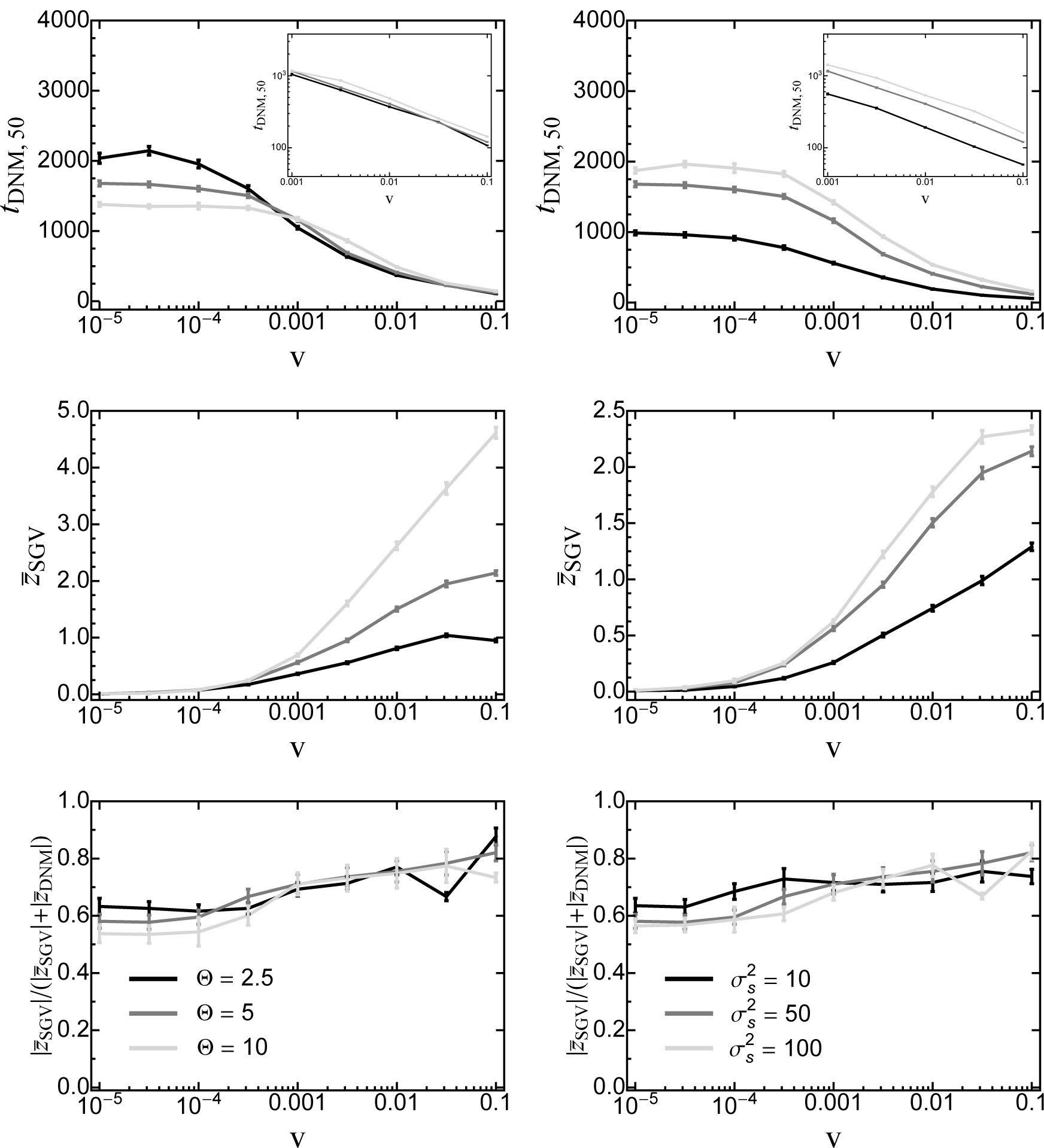
First row: the point in time *t*_DNM,50_ 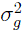 where 50% of the genetic variance is composed of *de-novo* mutations as a function of the rate of environmental change for various values of Θ (left) and 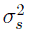 (right). Insets show the results for large *v* on a log-scale. Second row: The mean total phenotypic response from standing genetic variation at this time. Third row: The relative contribution of original standing genetic variation to the total genetic variance at time *t*_DNM,50_ 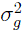. Data are means and standard errors from 1000 replicate simulation runs. Fixed parameters (if not stated otherwise): 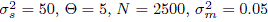.

